# Identification of a novel resistance gene which provides insight into Vip3Aa mode of action in *Helicoverpa armigera*

**DOI:** 10.1101/2024.08.11.607516

**Authors:** Andreas Bachler, Amanda Padovan, Craig J. Anderson, Yiyun Wei, Yidong Wu, Stephen Pearce, Sharon Downes, Bill James, Ashley Tessnow, Gregory A. Sword, Michelle Williams, Wee Tek Tay, Karl H. J. Gordon, Tom K. Walsh

## Abstract

The global reliance on *Bacillus thuringiensis* (Bt) proteins for controlling lepidopteran pests in cotton, corn, and soybean crops underscores the critical need to understand resistance mechanisms. Vip3Aa, one of the most widely deployed and currently effective Bt proteins in genetically modified crops, plays a pivotal role in pest management. This study identifies the molecular basis of Vip3Aa resistance in Australian *Helicoverpa armigera* through genetic crosses, and integrated genomic and transcriptomic analyses. We identified a previously uncharacterized gene, LOC110373801 (designated *HaVipR1*), as a crucial determinant of Vip3Aa resistance in two field-derived resistant lines. Functional validation using CRISPR-Cas9 knockout in susceptible lines confirmed the gene’s role in conferring resistance.

Despite extensive laboratory selection of Vip3Aa-resistant colonies in Lepidoptera, the biochemical mechanisms underlying resistance have remained elusive. Our research demonstrates that *HaVipR1*-mediated resistance operates independently of known resistance genes, including midgut-specific chitin synthase and the transcription factor SfMyb.

The identification of *HaVipR1* offers further insights into the Vip3Aa mechanism of action. This discovery is vital for devising strategies to counteract resistance and sustain the efficacy of Bt crops. Future research should focus on elucidating the biochemical pathways involving *HaVipR1* and investigating its interactions with other resistance mechanisms. Our findings underscore the utility of analysing field-derived resistant lines in providing biologically relevant insights and stress the necessity for comprehensive management strategies to maintain agricultural productivity.

**Significance Statement:** This is the first identification of a specific gene in *Helicoverpa armigera* which mediates resistance to the *Bacillus thuringiensis* (Bt) protein Vip3Aa. We identify that this gene is disrupted in two different ways in separate field-derived resistant lines, one being a large transposable element insertion in the first intron of the *HaVipR1* gene, confirmed using long-read sequencing. Disruption of this gene using CRISPR-Cas9 knockout in a susceptible *H. armigera* line confers Vip3A resistance. The identification of a specific gene is important for molecular monitoring and management of *H. armigera* as well as other global pests of concern like *Spodoptera frugiperda*. These findings also offer insights for researchers aiming to understand how Vip3Aa functions, as the action pathway remains unclear.

## Introduction

Proteins derived from *Bacillus thuringiensis* (Bt), primarily Cry toxins, have been used in crops to control lepidopteran pests since 1996 (Knight et al., 2021). In the last 20 years, a new type of toxin was discovered that is expressed during the vegetative growth phase of bacterial development. It is known as vegetative insecticidal protein (VIP) and a member of this group of toxins (Vip3A) has been added individually or in combination with Cry Bt proteins to genetically modified crops such as corn and cotton (Kurtz et al., 2007) (Estruch et al., 1996). This toxin is effective against a variety of lepidopteran pests and there appears to be no common mode of action between Vip proteins and Cry toxins (Ruiz de Escudero et al., 2014). In the USA and South America, large areas of corn and cotton with Vip3Aa have been planted since 2008 (International Service for the Acquisition of Agri-biotech, 2024; United States Environmental Protection, 2024). In Australia, Vip3A was deployed in 2016 as part of a triple stack together with Cry1Ac and Cry2Ab in cotton, primarily to control the global cotton bollworm, *Helicoverpa armigera* and an Australian local species *Helicoverpa punctigera*.

Despite this widespread use, the insecticidal mode of action of Vip3A in Lepidoptera has not been definitively established (Chakrabarty et al., 2020; Chakroun, Banyuls, Bel, et al., 2016). Understanding cases of resistance to Vip3A in previously susceptible insects can help shed light on the mechanisms of action. While there are some limited reports of field relevant Vip3Aa resistance and hints of resistance developing in some species, to date there have been no verified field failures from Vip3A resistance (Knight et al., 2021) (Yang, Kerns, et al., 2021) (Huang et al., 2023). As resources to better understand resistance, insect lines resistant to Vip3A have been isolated from field populations or selected in the laboratory, while its interaction withSf9 cell cultures from *Spodoptera frugiperda* has been investigated for potential insight into its mode of action and resistance mechanisms (Hou et al., 2021; Jiang et al., 2016). The development and maintenance of other Bt resistant lines of *H. armigera* in Australia has enabled the characterisation of resistance mechanisms. For example, the gene responsible for Cry2Ab resistance was identified from a field isolated resistant line (Tay et al., 2015). Similarly, the detection of resistant alleles and the subsequent establishment of Vip3A resistant lines has allowed the characterisation of the resistance phenotype (Joaquin Gomis-Cebolla et al., 2018; Mahon et al., 2012).

A recent review summarised the current knowledge about the mode of action and resistance mechanisms to Bt toxins including Vip3A (Jurat-Fuentes et al., 2021). Briefly, ingested, full length Vip3Aa is thought to be activated by proteases before binding and forming pores in brush border membrane vesicles (BBMVs), dissected midguts, and planar lipid bilayers (Lee et al., 2003) (Liu et al., 2011). This pore formation then causes extensive damage to the midgut, with lysed and swollen cells leaking cellular material to the gut lumen (Gomis-Cebolla et al., 2017). An alternative mechanism has also been proposed where Vip3Aa and Vip3C exposure triggers midgut cell apoptosis in *S. exigua* larvae (Hernández-Martínez et al., 2017), and Vip3Aa causes the same in cultured Sf9 cells (Jiang et al., 2016).

Binding sites for Vip3A toxins appear to be generally distinct to Cry1 and Cry2 type toxins (Kahn et al., 2018), except possibly for Cry1Ia in *Spodoptera eridania* (Bergamasco et al., 2013). This observation explains lack of relevant cross-resistance to Vip3 in Cry1-or Cry2-resistant lepidopteran colonies (Tabashnik & Carrière, 2020). Furthermore, work on dual resistant individuals has shown that the two resistant phenotypes are not linked in *H. armigera* and *H. punctigera* (Walsh et al., 2014). There is no evidence of cross resistance between classes of Bt proteins (ie: Cry1 type, Cry2 type and Vip type proteins) in these mechanisms suggesting independent modes of action and resistance for these toxins in *H. armigera* (Joaquin Gomis-Cebolla et al., 2018). However, within the Vip family of proteins there was cross resistance between Vip3Aa and Vip3C, which share less than 45% protein sequence identity (Gupta et al., 2021). Vip3A and Vip3C proteins share binding sites as shown through competitive binding assays, suggesting that a binding site alteration could confer resistance to these two families of Vip3 proteins (Kahn et al., 2018) and this cross resistance has been observed in *H. armigera* (J. Gomis-Cebolla et al., 2018). However as with the Cry1 type Bt toxins, it is possible that Vip3 proteins interact with more than one class of proteins in the insect midgut (Jurat-Fuentes et al., 2021).

Resistance to Vip3Aa has already been identified in at least five Heliothine species. Even before commercialization of Vip3A, Vip3A-resistant alleles were detected in Australian *H. armigera* (at 2.7% frequency) and *H. punctigera* (0.8%) using F_2_ screens (Mahon et al., 2012) and these frequencies have remained more or less the same between 2009 and 2023 (Knight et al., 2021). F_2_ screens for major Vip3Aa resistance alleles in *H. zea* from Texas (United States) have estimated 0.65% frequency (Yang et al., 2020), with another study from Louisiana (United States) having an estimated frequency of 1.49% (Lin et al., 2022). An F_2_ screen for Vip3Aa resistance in *S. frugiperda* from Brazil indicated a 0.09% overall frequency of resistance alleles (Bernardi et al., 2015). Similar screens in Louisiana and Florida (United States) resulted in an estimated resistance allele frequency of 0.48% in *S. frugiperda* (F Yang et al., 2013; Yang et al., 2018). Selection for Vip3Aa resistance in *Heliothis virescens* over 13 generations resulted in insects with a resistance ratio of 2,040 fold relative to susceptible insects (Pickett et al., 2017). Similar laboratory selections for Vip resistance has resulted in high resistance ratios in many species, including *Spodoptera frugiperda* with > 9,800-fold resistance (Wen et al., 2023) and >3200-fold resistance (Bernardi et al., 2016) as well as in *H. zea* with >588-fold resistance (Yang et al., 2020). The majority of Vip3A resistance in Lepidoptera has been identified as recessive and likely conferred by a single autosomal gene with no cross-resistance to Cry1Ac or Cry2Ab (Mahon et al., 2012) (Joaquín Gomis-Cebolla et al., 2018) (Yang, Santiago González, et al., 2021). Recent investigations have identified potential selection for multi-genic Vip3A tolerance in in *Helicoverpa zea* (Pezzini et al., 2024). However, the underlying genetic mechanism of Vip resistance is unclear in most species, with the first report of a chitin synthase gene being involved in Vip3A resistance in *S. frugiperda* recently (Liu et al., 2024).

*Summary* The first (Pearce et al. 2017) and subsequent (Zhang et al., 2022) *H. armigera* genomes have opened up new avenues of research into this costly pest including the modes of action and mechanisms of resistance to Bt proteins. In this work we describe the characterisation of a Vip3A resistant line (“Ha85”, previously identified as SP85 by (Chakroun et al., 2016)), conducted genetic mapping of Vip3A resistance, and identified a candidate locus. Using transcriptome sequencing, we pinpointed the gene responsible for Vip3A resistance. We confirmed the role of this gene by employing CRISPR-Cas9 to knock it out, successfully generating a resistant phenotype. We also explored the potential function of this gene in Vip3A resistance and examined another Vip3A-resistant line (Ha477) identified from the field through resistance monitoring. Using long-read sequencing approaches, we discovered significant variation in the same Vip3A resistance gene.

## Results

### Genomic mapping of resistance loci

To identify genomic regions under selection for Vip3A resistance, progeny from a female-informative backcross were analysed using Restriction site-associated DNA sequencing (RAD-seq). Progeny were either exposed to a Vip3A surface overlay (selected) or left untreated (control). A total of 32,920 loci were identified in the treated (n=20) and control (n=20) populations using STACKS analysis with the HaSCD2 reference. Smoothed F_st_ and Phi (Φ_st_) were assessed for all loci. Invalid values (such as those with negative Fst) were set to 0. All chromosomes had between 400 and 2,000 valid loci able to be assessed (see Table S1 for raw counts). The mean value for F_st_ and Φ_st_ was assessed in 1 mega-base pair (mbp) genomic windows for each chromosome. The results indicated a single chromosome (chromosome 2) under selection for Vip3A resistance in the selected population (Fig. 1).

**Figure 1.**
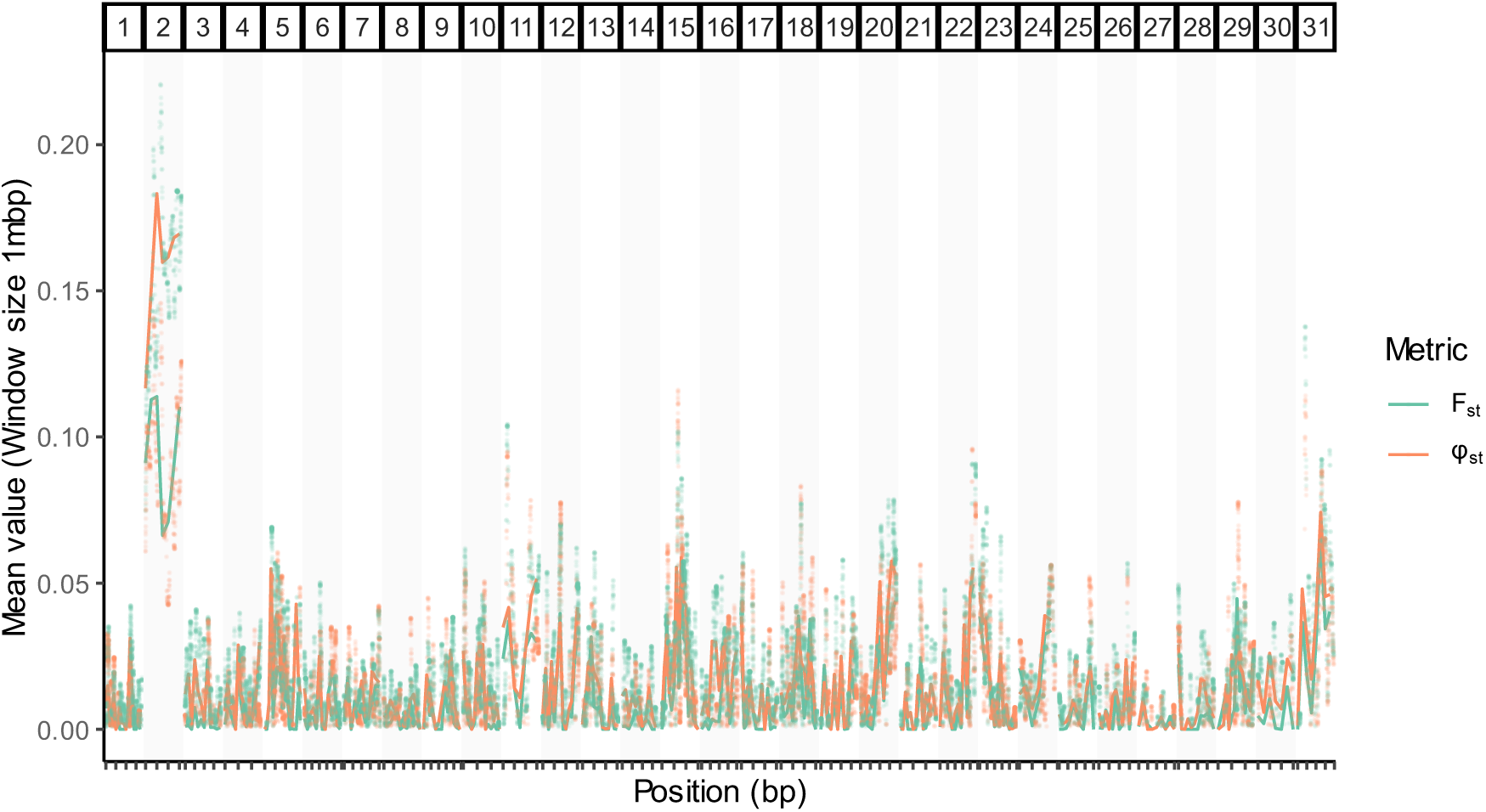
Mapping of resistance associated loci from a female-informative cross indicates a single chromosome as under selection for Vip3A resistance. F_st_ and Phi (Φ_st_) values were derived from RAD-Seq data to identify genomic regions under selection for Vip3A resistance in progeny from a female-informative backcross. Genetic markers across all chromosomes were assessed using STACKS, with raw points shown above and the mean of F_st_ and Φ_st_ metrics shown in 1mbp genomic windows per chromosome. Progeny were either exposed to a Vip3A surface overlay (selected) or left untreated (control).

### Transcriptomic analysis of Vip3A resistant lines

Differential gene expression (DGE) between Vip3A-resistant (Ha85) and susceptible (GR) lines was assessed using pooled RNA from mid-gut tissues. Analysis was performed with DeSeq2, and genes with a total combined count of less than 500 reads were omitted, resulting in the assessment of 5,663 genes. The results are visualized in a Manhattan plot, highlighting some differential expression across the 31 *H. armigera* chromosomes, with the most significant gene being LOC110373801 on chromosome 2, annotated as “Uncharacterized protein.” (Fig 2. A). We refer to this as *HaVipR1* throughout this manuscript. The expression of this gene was almost 10-fold lower in the resistant vs the susceptible population (Fig 2. B). The differential expression observed in RNA-Seq was validated using qPCR analysis on mid-gut samples from the Vip3A-resistant (Ha85), susceptible (GR), and an additional resistant line (Ha477, allelic for resistance to Ha85). The transcription levels of LOC110373801 were significantly down-regulated in both resistant lines compared to the susceptible line, corroborating the DGE results (Fig 2. C).

**Figure 2.**
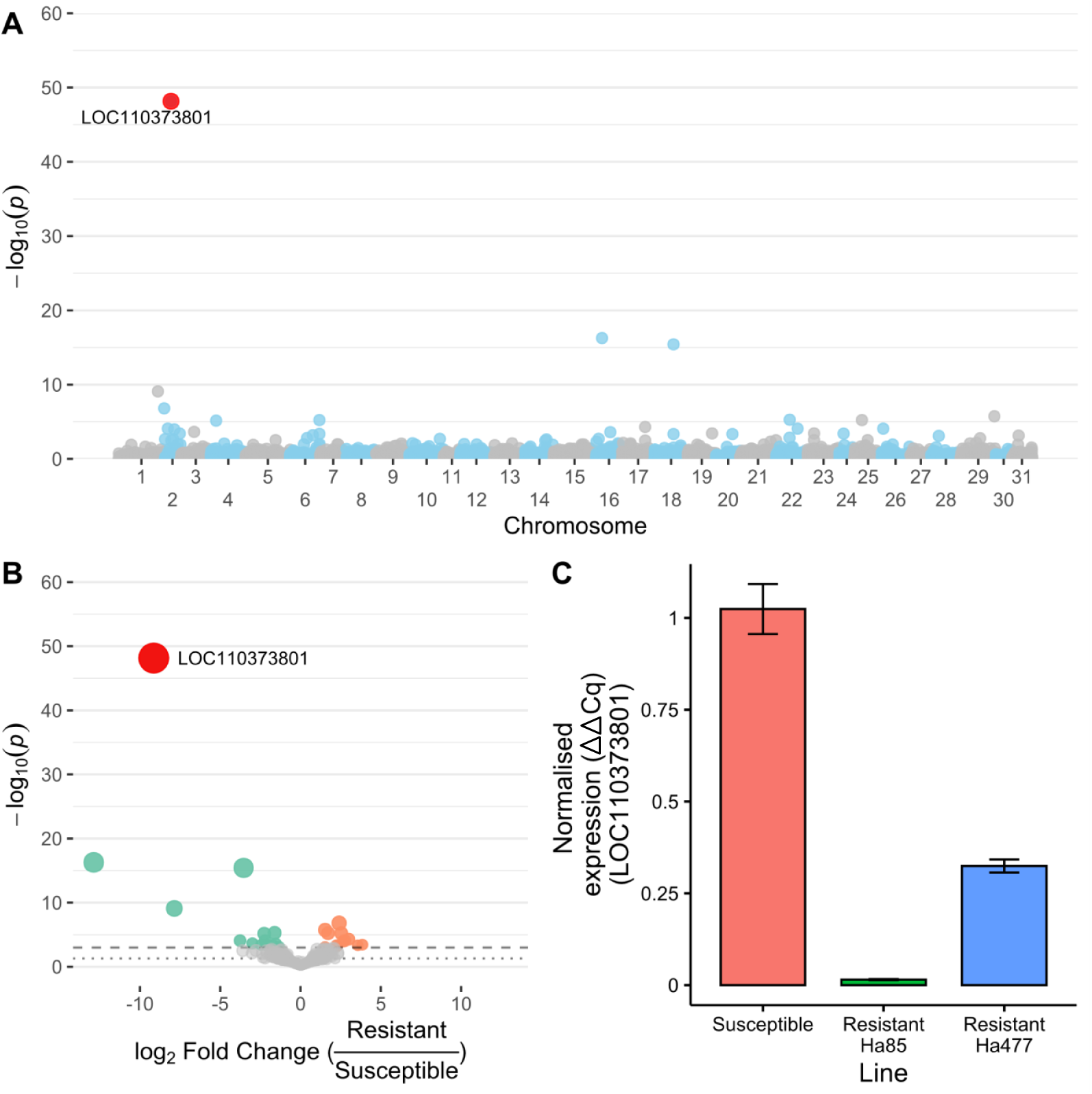
Differential gene expression in Vip3A-resistant versus susceptible Helicoverpa armigera lines. A) Pooled RNA mid-gut tissues from a resistant (Ha85) and susceptible (GR) line were analysed for differential gene expression analysis with DeSeq2. A Manhattan plot of assessed genes was generated for all assessed genes across the 31 H. armigera chromosomes. The gene exhibiting the most significant differential expression (LOC110373801, annotated as “Uncharacterised protein”) is highlighted. B) Visualization of differential expression between the resistant and susceptible lines using a volcano plot. Significance thresholds are indicated by dashed (p < 0.01) and dotted (p < 0.05) lines. C) Quantitative PCR (qPCR) analysis of the transcription level of LOC110373801 in mid-gut samples from three distinct groups of H. armigera: the same Vip3A-resistant (Ha85) and susceptible (GR) lines used in the transcriptome analysis, and an additional Vip3A-resistant line (Ha477), allelic for resistance to Ha85. Transcription levels were normalized to the reference gene EF-1α.

### Generation and validation of a *HaVipR1* knockout strain using CRISPR

Fresh eggs were injected individually with one nanolitre mix of the Cas9 protein and the sgRNA of *HaVipR1.* Among the injected eggs, 14.6% (135/925) of them hatched. Among the 135 neonates, 57.8% (78/135) of them developed into adults (G0). The G0 moths were individually crossed with SCD moths (single-pair mating) to produce the next generation (G1). Forty-eight single pair families produced fertile progeny. Six to 10 larvae were randomly selected from each single-pair family to check if there were inherited indel mutations, and 30 out of the 48 single pair families had at least one indel mutation created by Cas9 using PCR amplification. When a cluster of double peaks was observed around the Cas9 induced DSB cutting site (Fig 3), PCR products were TA-cloned and sequenced by Life Technology (Shanghai, China) to determine the exact indel mutation type. Most abundant indel mutations flanking the target site were listed in Figure 4 for the G_1_ larvae from the single-pair families (G_0_ × SCD).

**Figure 3.**
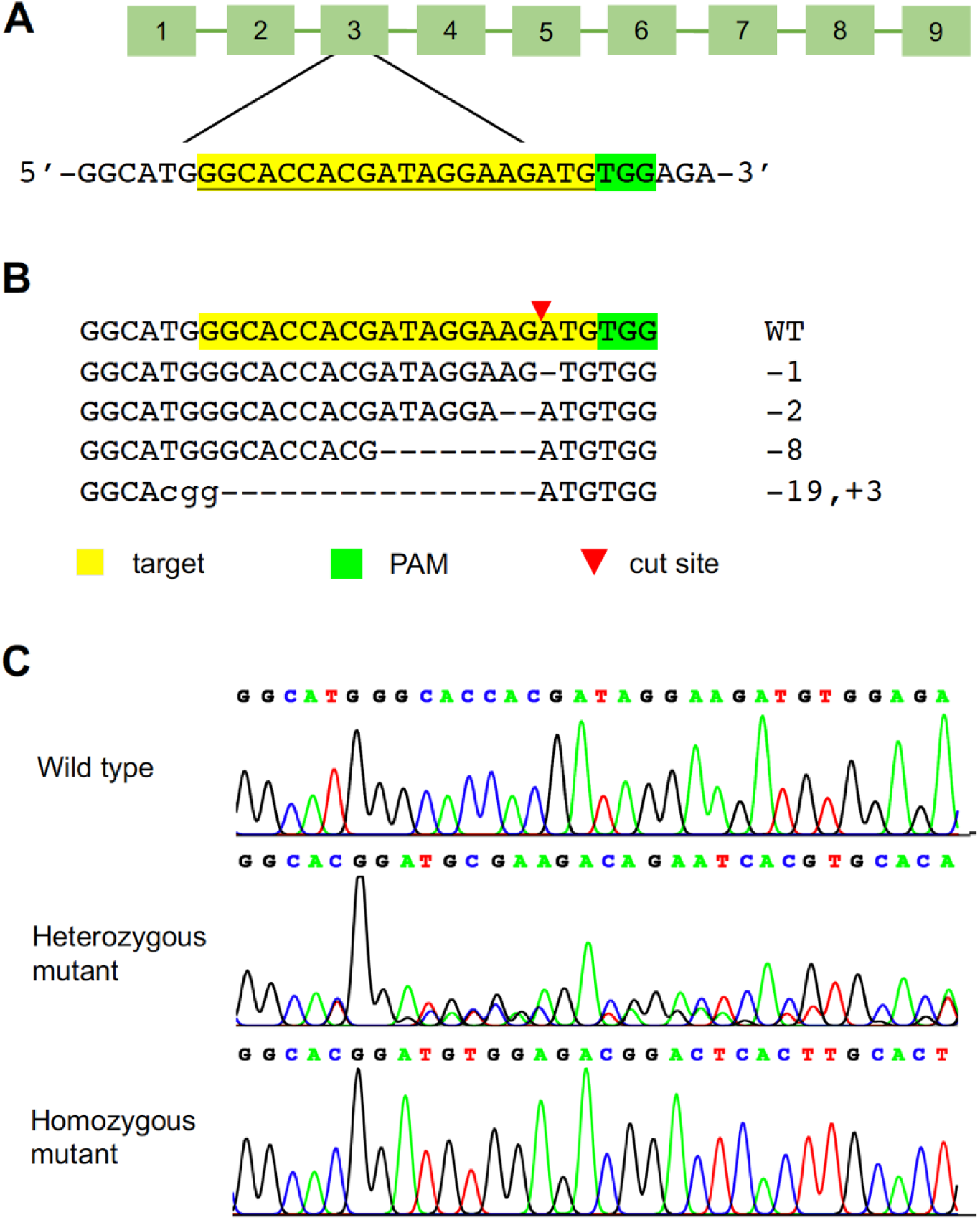
(A). Schematic diagram of the sgRNA-targeted site and sequences in the exon 3 of HaVip3R1. (B). Sequences of indel mutations flanking the target site in the G1 larvae from the single-pair families between G0 and SCD. The target sequences of the wild type HaVip3R1 allele are highlighted in yellow and the PAM sequences in green. The cleavage site is indicated by a red triangle. Deleted bases are represented by dashes and inserted bases are shown in lower case. The numbers of bases deleted or inserted are listed at the right side of sequences. (C). Representative chromatograms of direct sequencing of PCR products for genotyping the 16-bp deletion of HaVip3R1 created by CRISPR/Cas9.

**Figure 4.**
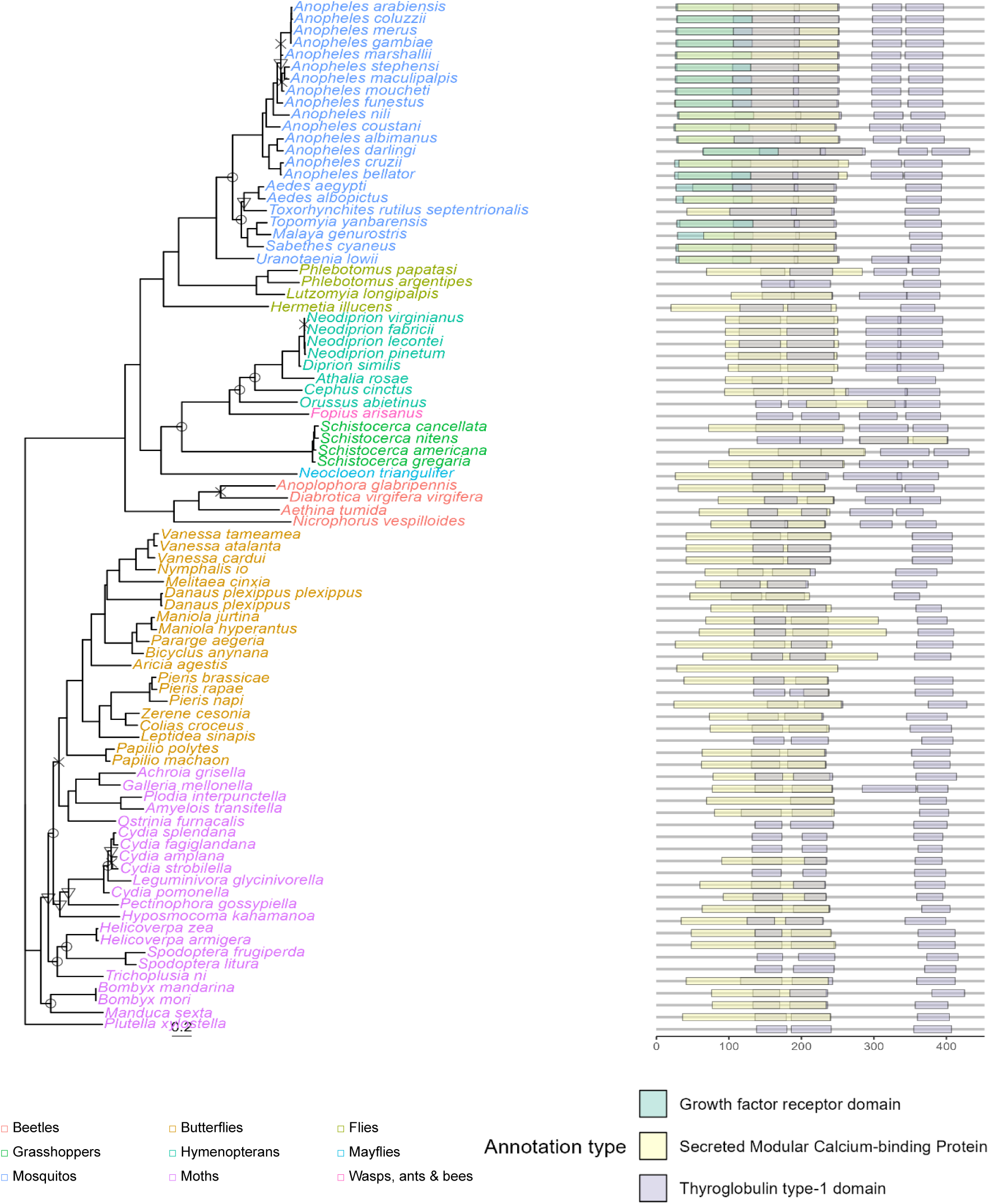
Phylogenetic conservation and functional domain analysis of the HaVipR1/LOC110373801 gene across diverse insect taxa. A) Phylogenetic analysis of the HaVipR1 gene conducted using BlastP to identify top protein hits, followed by sequence alignment with MUSCLE. The phylogenetic tree was constructed using IQ-TREE, employing ModelFinder for model selection and 1000 Ultrafast bootstraps to assess node support. Bootstrap support levels are denoted as follows: less than 70% (✕), 70–80% (▽), and 80–90% (○). Nodes with support greater than 90% are not displayed. B) Assessment of functional domains within the protein sequences using InterProScan. The results, including annotations from SUPERFAMILY and PANTHER databases, are presented for each protein in the phylogeny.

Fifty-three pupae from one of 30 single pair families were selected for genotyping at exon 3 of *HaVipR1* using exuviates of the final instar larvae (as in Wang et al., 2016). Among the 53 individuals sequenced, 20 were heterozygous mutants of 2-bp deletion, 14 were heterozygous mutants of 16-bp deletion, 7 were heterozygous mutants of 3-bp deletion, 1 was heterozygous mutants of 1-bp deletion, and 11 were wild type homozygotes. To transmit the mutant allele to G2, we mass crossed the 6 male moths and 8 female moths which were heterozygous for the 16-bp deletion in exon 3 of *HaVipR1*.

Neonates of G2 were screened for resistance with a discriminating concentration of 50 µg Vip3Aa/cm2, and 19.1% survived (78/408). Pupae of the 48 survivors and 52 untreated larvae were genotyped for the 16-bp deletion in exon 3 of *HaVipR1* using exuviates of the final instar larvae. For the untreated 52 larvae, the expected 1:2:1 ratio was observed: homozygous wild type (12), heterozygotes (28), and homozygous mutant type (12). In contrast, all of the 48 survivors were identified as homozygous for the 16-bp deletion exon 3 of *HaVip3R1*. To statistically confirm this association, we performed a chi-square test of independence (degrees of freedom = 2). The observed and expected genotype frequencies are summarized in Table 1. The chi-square test resulted in a χ² value of 61.538 and corresponding p-value of 4.336e-14, which is smaller than the threshold for significance of 0.05, rejecting the null hypothesis of independence. These results provide strong statistical evidence of a significant association between the Vip3Aa resistance phenotype and the 16-bp deletion mutation of *HaVipR1*. The 48 survivors were pooled and mass crossed to generate a Vip3Aa-resistant strain, named 548KO2.

**Table 1.** Observed genotyping results following selection of CRISPR disrupted mutants (*r* indicates the 16-bp deletion, *s* indicates the susceptible wild-type).

|  | *ss* | *rs* | *rr* | *Total* |
| --- | --- | --- | --- | --- |
| Survivors (48) | 0 | 0 | 48 | 48 |
| Control (52) | 12 | 28 | 12 | 52 |
| Total | 12 | 28 | 60 | 100 |

### Phylogenetic and functional analysis of *HaVipR1* gene

#### Phylogenetic conservation

The protein product of the *HaVipR1* gene (LOC110373801) was analysed for phylogenetic conservation across diverse insect taxa. A phylogenetic tree was constructed using sequences identified via BlastP and analysed with IQ-TREE (Figure 4). The generated phylogenetic tree revealed that the *HaVipR1* gene is relatively conserved within moth and butterfly groups, forming a distinct clade separate from other insects, such as beetles and mosquitoes, which showed higher divergence in sequence homology and functional annotations.

#### Functional annotation

Functional domains within the *HaVipR1* protein sequences were assessed using InterProScan and annotations from the SUPERFAMILY and PANTHER databases were included for each sequence and indicated the presence of conserved thyroglobulin type-1 (SSF57610/IPR036857) and protease inhibition (PTHR12352 / IPR051950) domains across almost all sequences. The presence of these domains provides significant and novel insight into the mode of action of the Vip3A toxin protein (see Discussion). The mosquito clade included an additional domain relating to growth factor receptor domain (SSF57184/IPR009030). PANTHER GO terms indicated involvement in the biological process of cell-matrix adhesion (GO:0007160), with cellular components including the basement membrane (GO:0005604) and extracellular space (GO:0005615), while no specific molecular functions were identified.

### Structural variation in resistant and susceptible lines

Genetic analysis of the *HaVipR1* gene in Vip3A-resistant and susceptible *H. armigera* lines revealed significant structural variation (Figure 5). In the resistant Ha85 line, a 149 bp deletion in the 5’ untranslated region (5’ UTR) was identified, removing the Kozak sequence and impacting gene expression. In the resistant Ha477 line, a large transposable element was found, extending intron 1 by ∼10 kbp and impacting gene splicing, with spliced alignments identified near the 3’ end of the repeat-rich region to exon 2 of the *HaVipR1* gene (see Fig. S1).

**Figure 5.**
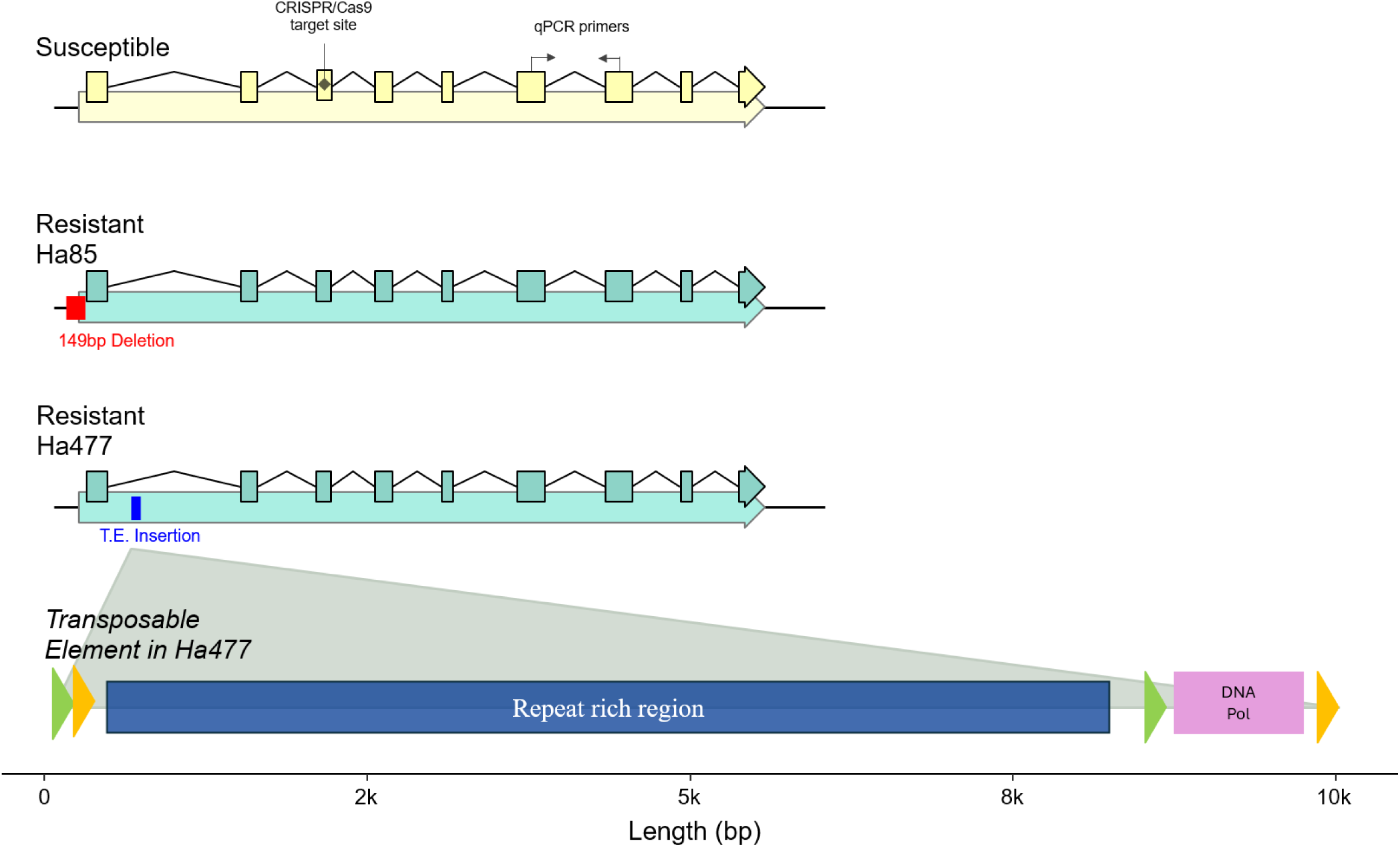
Genetic variation of the HaVipR1 gene in susceptible and resistant Helicoverpa armigera lines. The CRISPR/Cas9 cut sites along with the locations of qPCR primers are marked on the gene model of the susceptible line. A 149 bp deletion in the 5’ untranslated region (UTR) of the resistant Ha85 line is highlighted with a red box, indicating the genetic alteration contributing to resistance. The insertion site of a transposable element in the resistant Ha477 line is marked with a blue box, showing the site of genomic integration. Detailed representation of the large transposable element features two distinct regions: a large repeat-rich area (coloured in dark blue) and a region with high sequence similarity to a DNA polymerase. Repeat elements and orientation are indicated with orange and green arrows. The transposable element and gene models are drawn to scale.

## Discussion

Globally, the control of lepidopteran pests in cotton, corn, and soybean heavily relies on Bt proteins, with increasing prevalence of Vip Bt toxins (Tabashnik et al., 2023). In this study, we identified the *HaVipR1* gene (LOC110373801) as being strongly associated with Vip3Aa resistance in two field-derived resistant lines of *H. armigera*. This identification was achieved through a combination of genetic crosses, genomic, and transcriptomic analyses. We then used CRISPR to knock this gene out in susceptible *H. armigera* lines conferring resistance and confirming a role for this gene in Vip3A resistance.

This gene appears to be distinct from currently known Vip3A associated genes. In *Spodoptera frugiperda*, two candidate Vip3A resistance genes have recently been reported (the transcription factor “SfMyb” (Jin et al., 2023) and the chitin-synthase gene “SfCHS2” (Liu et al., 2024)). The domains identified in the *HaVipR1* protein show it does not have a similar function to either of these two proteins (see Table S3). Analysis of the homologues of the *S. frugiperda* Vip3A resistance genes in the susceptible and resistant *H. armigera* transcriptome data also shows no statistically significant association with Vip3A resistance (Fig S2). Further assessment of *HaVipR1* in relation to other known Vip3A receptor genes, generally studied in Sf9 cell lines, revealed no significant association in the resistant lines (Fig S2). These receptor genes include ribosome S2 protein 3, tenascin-like glycoprotein 4, fibroblast growth factor receptor-like protein 5, and scavenger receptor class C-like protein. The expression of the *HaVipR1* gene appears limited to the mid-gut and to be extra-cellularly located (assessed in both *H. armigera* (Fig. S3 & Fig S4) and for *Bombyx mori* (Fig S5). *HaVipR1* does not appear to be expressed in Sf9 cell lines, which are ovarian derived (Fig S6). These findings suggest that *HaVipR1* represents a novel mechanism of Vip3Aa resistance, distinct from previously identified genes and receptors.

The function of HaVipR1, and mechanism by which its absence leads to Vip3A resistance, is difficult to discern. The gene is widespread across insects and in Lepidoptera generally contains two thyroglobulin domains and a calcium-binding domain (see Fig 4). The disruption of this gene in the resistant line appears to have some fitness cost in the absence of selection (see Figure S7). The thyroglobulin domain has been understood to function as a potent cysteine protease inhibitor, specifically of Cathepsins B and L (Mihelič & Turk, 2007) (Turk et al., 2012). Activation of Vip3A in the mid-gut requires processing from the inactivated proto-toxin form to the activated toxin by mid-gut proteases (M. Chakroun et al., 2016). Disruption of *HaVipR1* could leads to lower inhibition of the proteases, and subsequent higher cysteine protease activity in the midgut. This may lead to more rapid degradation of activated Vip3A, enabling resistance. However, previous work characterising the resistant strain analysed here identified the opposite case, where the mid-gut juices from the resistant line had a decreased proteolytic processing of the proto-toxin to the activated toxin, and after 60 minutes both the resistant and susceptible strain had identical levels of activated Vip3A toxin (Maissa Chakroun et al., 2016). In addition, proteases that activate Vip3A are generally serine proteases, not cysteine proteases, and generally serine proteases have been identified to be differentially expressed in response to Vip3A treatment (Bel et al., 2013). The identified domains of *HaVipR1* limit its potential role in directly mediating proteolytic activity of proteases on Vip3A, however further investigations to conclusively exclude this mechanism are required.

Alternatively, *HaVipR1* may have an undescribed function linked to a receptor that mediates Vip3A binding. Changes in receptor binding are highly relevant in mediating resistance in Cry Bt toxins (Pigott & Ellar, 2007) and associated with Vip3A resistance in *Helicoverpa zea* (Kerns et al., 2023). However, analysis of BBMV derived from the resistant and susceptible lines failed to identify any statistically significant differences in the binding of Vip3A (Maissa Chakroun et al., 2016). While changes were not seen in binding to BBMVs derived from the resistant line, a reduction in the peritrophic membrane were observed in the case of the chitin-synthase Vip3A resistance in *S. frugiperda*, indicating a potential interaction of Vip3A with other areas of the mid-gut epithelium, or other mid-gut cell types. The presence of *HaVipR1* in the extra-cellular space may indicate a potential interaction with the peritrophic membrane, and changes in permeabilization of the peritrophic matrix (PM) have been indicated to be associated with other types of pathogen resistance in *H. armigera* (Jakubowska et al., 2010).

A novel mechanism for resistance could be indicated by the presence of both the thyroglobulin and calcium binding domains of *HaVipR1*, implicating a potential role in mediating repair of pores formed by activated Vip3A. Cysteine proteases, specifically cathepsin B and L are involved in targeting repair of pores in cell culture in human cell lines(Castro-Gomes et al., 2016). In addition, calcium ions are key signal for initiating plasma membrane repair (McNeil & Kirchhausen, 2005). In its native role, *HaVipR1* may act to regulate the degree of pore repair occurring for damaged cells, helping balance the turnover of mid-gut cells between apoptosis and autophagy (Turk et al., 2012; Vidak et al., 2019). Disruption of *HaVipR1* gene expression may lead to increased repair of pores generated by active Vip3A, and potentially shifts the balance of mid-gut cells from apoptosis to autophagy (Saikhedkar et al., 2015). In this context it is noteworthy that the Myb transcription factor, knockout of whose expression in *S. frugiperda* confers resistance to Vip3A (Jin et al., 2023), drives expression of Bcl2 (Ciciro & Sala, 2021), an apoptosis inhibitor which also affects autophagy (Levine & Kroemer, 2019). While this mechanism encompasses the currently described domains of *HaVipR1* it struggles to explain the lack of cross-resistance observed, as increased pore repair would be a general mechanism of Bt toxin resistance. The specificity of this resistance mechanism may be explained by differences in ion selectivity between Cry and Vip pores (Lee et al., 2003). However, further work exploring the cellular location of this gene, and any potential interaction partners, will be required before this potential mechanism of action can be validated, although some indications exist that repair mechanisms may be involved in Bt resistance (Castagnola & Jurat-Fuentes, 2016). Further research into this area will be crucial for understanding of the mechanism of action of Vip3A.

The two identified resistant lines, Ha85 and Ha477, exhibit distinct genetic alterations leading to the disruption of the *HaVipR1* gene. In Ha85, a small deletion immediately upstream of the start codon disrupts gene function, while Ha477 harbors a large transposable element insertion within the first intron sequence, which disrupts regular splicing of this gene (see Fig S1). Transposable elements have been increasingly implicated in resistance to various Bt toxins, including Cry1Ac (Gahan et al., 2001), Cry2Ab (Guo et al., 2023), and more recently, Vip3A (Liu et al., 2024). The transposable element identified in Ha477 is the largest reported so far. Its classification remains unclear due to the absence of recognized open reading frames that match previously identified elements. It is potentially a composite transposable element, comprising a Class II DNA transposon with an additional insertion of a Class I Retrotransposon (Wells & Feschotte, 2020). The large size and complex structure of this transposable element pose significant challenges for identification using short-read sequencing technologies. The repetitive regions hinder effective short-read mapping, emphasizing the need for long-read sequencing to accurately characterize such elements. This has important implications for molecular field monitoring applications, where long-read sequencing would be crucial for identifying resistance alleles in natural populations. The identification of these transposable element insertions underscores the complexity of resistance mechanisms involving transposable elements and highlights the necessity for the use of the latest genomic tools in resistance monitoring and management. The information that transposable elements in introns of genes can dysregulate function provides further evidence of the importance of identification of transposable elements in Vip3A resistance, and will aid in developing more effective monitoring strategies for combating resistance and ensuring the sustainability of Bt crops.

In summary, our study highlights the identification of *HaVipR1* as a key gene in Vip3Aa resistance in *H. armigera* and provides insights into the potential mechanisms of resistance. The involvement of transposable elements further complicates the resistance landscape and underscores the necessity for advanced sequencing technologies in resistance monitoring and management. Understanding these mechanisms is crucial for developing effective strategies to combat resistance and ensure the sustainability of Bt crops.

## Materials and Methods

### Vip3A Protein Expression

The Vip3A protein utilized in this study originated from three sources. The variant for fitness, monitoring, and bioassay tasks was a Vip3Aa clone provided under a Material Transfer Agreement by Bayer/BASF. For resistance allele identification and subsequent characterization, Vip3Aa sequences were isolated from *Bacillus thuringiensis* isolates maintained at the CSIRO Black Mountain Innovation Precinct. The genomes of these isolates were sequenced using an Illumina MiSeq system and assembled via CLC Genomics software. Vip3Aa sequences were identified using BLAST, with specific primers designed to amplify these sequences. Amplified PCR products were cloned into the pGEX4T-1 vector and expressed in *Escherichia coli*. The Vip3Aa used in validation of the CRISPR-knockout was provided by the Institute of Plant Protection, Chinese Academy of Agricultural Sciences (CAAS), Beijing, China.

### Insect Lines, Rearing, and Bioassay

Insect lines used for linkage mapping, fitness testing, and resistance monitoring, specifically HA85 and Ha477, are Vip3A-resistant *H. armigera* lines derived from field populations. These lines have been detailed in previous publications (Chakroun et al., 2016) and have been maintained and reselected in a laboratory in Canberra, Australia since 2010. Both lines were backcrossed into the susceptible GR line at least five times, achieving an approximate 92% allelic similarity with the susceptible line. Bioassays to test resistance were conducted using a discriminating dose of 10 µg/cm^2^ of Vip3A, applying the toxin to the surface of the diet (approach based on (Akhurst et al., 2003)). Outcomes were evaluated based on mortality and development after 7 days.

To assess the fitness cost associated with Vip3A resistance, a population cage experiment was conducted, following a protocol similar to (Mahon & Young, 2010). The initial experimental setup included 60 individuals from each population: the Vip3A-resistant population (HA85) and the susceptible population (GR). These populations were mixed to achieve an initial allele frequency of 50% and were maintained in two replicates for 8 generations.

Both populations were not subjected to Vip3A selection pressure throughout the experiment. Over eight generations, a subset of individuals from each replicate population was phenotypically screened using the same Vip3A discriminating dose as described above. The proportion of Vip3A resistance is shown for each generation in Supplementary Figure S7.

### Linkage mapping of resistance allele

#### Female Informative Cross

Using the absence of recombination during meiosis in female Lepidoptera, the Vip3A resistant locus was mapped to a specific linkage group. This was achieved using a cross involving an HA85 family (family F2031; HA85 Vip3A R+/R+♂ x GR Vip3A R-/R-♀). Individual F_1_ females were subsequently backcrossed to resistant HA85 males, with offspring screened for resistance using a discriminating dose of Vip3A (10 µg/cm^2^). For control, over 100 F_2_ offspring were not bio-assayed, but were included in subsequent genotyping efforts.

#### DNA Extraction and RADseq Analysis

Parents from both crosses and selected and control offspring were preserved in 100% ethanol at −20°C. DNA extraction was performed using the Qiagen Blood and Tissue Kit (Qiagen, Netherlands) following the manufacturer’s instructions. RADseq (Restriction Associated DNA Sequencing) libraries were generated for the female informative cross, following the protocol outlined by Etter et al. (2011), utilizing PstI as the restriction enzyme. RADtag libraries were sequenced on an Illumina HiSeq platform at the Biological Resources Facility (BRF), Australian National University, Canberra, Australia. Resulting sequences were processed using STACKS software (v2.59)(Catchen et al., 2013), aligned to the chromosomal genome (Ha2SCD, GCF_023701775.1), and population statistics plotted for valid loci in the selected and control populations for the female informative cross.

### RNA sequencing and transcriptome analysis of resistant lines

RNA was isolated from midguts of *Helicoverpa armigera* using Trizol reagent (Life Sciences, Carlsbad, CA, USA) according to the manufacturer’s instructions. The initial transcriptomic analysis involved RNA extracted from a single replicate of five pooled midgut individuals from both the resistant (HA85) and susceptible colonies, which was sequenced at BGI, Shenzhen, China. A subsequent follow-up analysis involved three replicates of five pooled midguts from Ha85, Ha477 and the susceptible strain, GR, also sequenced at BGI. Transcript mapping was performed against the chromosomal *H. armigera* genome (HaSCD2, GCF_023701775.1) using Hisat2 (v2.2.1 (Langmead & Salzberg, 2012), and differential expression analysis was conducted for the Ha85 and susceptible line using DESeq2 (v1.44.0) (Love et al., 2014). Read counts and proportion of reads mapping are provided in Table S2. Raw sequence data are available under BioProject PRJNA1119665.

### qPCR confirmation of down-regulated *HaVipR1* gene

Primers were designed to target the differentially regulated candidate gene identified in the RNAseq analysis. Association of the resistance allele with the candidate transcript was evaluated through PCR in various samples, including individuals from the HA85 mapping family, the Ha477 resistant colony, a double-resistant line (”DRES”) derived from Ha477 and SP15 (a Cry2Ab-resistant line), and families identified as resistant in the Vip3A resistance monitoring project. For the qPCR analysis, midguts from late 3rd instar larvae of *H. armigera* lines GR, Ha85, and Ha477 were dissected and snap-frozen in liquid nitrogen. RNA extraction from the midguts was performed using the RNeasy Mini Kit (QIAGEN, Hilden, Germany, Catalog No. 74104). We employed the Luna Universal One-Step RT-qPCR Kit for the qPCR reactions (New England Biolabs, Ipswich, MA, USA, Catalog No. E3005). Each 20 μL reaction consisted of 10 μL reaction mix, 1 μL enzyme mix, 1.6 μL each of forward and reverse primers, 2 μL RNA (diluted to 150 ng/μL), and 5.4 μL water. The reference gene EF-α was used, with five biological replicates and three technical replicates for each gene. Primer sequences were verified by PCR and Sanger sequencing using ‘Big Dye’ chemistry. The primer sequences used were as follows:

Vip3A primers:

VipF2: 5’-GGA AGG GTA CAA CGT GGA CT-3’
VipR2: 5’-CAC GCT CGT CGA TAC AGA TT-3’

Reference gene primers:

EF1α Forward: 5’-GAC AAA CGT ACC ATC GAG AAG-3’
EF1α Reverse: 5’-GAT ACC AGC CTC GAA CTC AC-3’

All qPCR reactions and data analysis were conducted on a BioRad real-time PCR system instrument.

### Long-read sequencing and *de novo* assembly

High-molecular-weight (HMW) DNA was extracted from the head and thorax (approximately 50 mg of tissue) of an adult moth from the Ha477 line using the Circulomics HMW Nanobind kit (Circulomics, Catalog No. 102-301-900). DNA extraction quality and quantity were assessed using the Promega QuantiFluor dsDNA system. The extracted DNA was then submitted to the Australian Genome Research Facility (AGRF) for long-read sequencing using Oxford Nanopore Technologies (ONT). The HMW DNA sample was processed for sequencing by AGRF following the PromethION P2 Kit V13 (SQK-LSK113) barcoding protocol and sequenced on a PromethION flow cell (type R10.4.1, FLO-PRO114M). Base-calling was performed directly on the flow cell using High-Accuracy mode. Additionally, a matched Illumina short-read library was generated on a NovaSeq S4. The raw data have been deposited in the SRA under accession number PRJNA1119665.

The long-read data was *de novo* assembled using Flye (v2.9.3) (Kolmogorov et al., 2019) and polished using the matched short-read data for three rounds with Pilon (v1.24) (Walker et al., 2014), following the recommended protocol. The resulting assemblies were then mapped to the reference using minimap2 (v2.25) (Li, 2018), sorted and indexed using samtools (v1.18) (Danecek et al., 2021). The region containing the *HaVipR1* gene was extracted from the polished assembly, and the repeat content was analyzed using the Geneious Prime (2023.2.1) Repeat Finder tool. RNA-seq data from the susceptible and Ha477 lines were aligned to the polished Ha477 assembly using the same alignment protocol as for transcriptomic analysis, and splicing was assessed visually using the Integrated Genomics Viewer (IGV, v 2.17).

### Phylogenetic analysis of the *HaVipR1* gene

The *HaVipR1* gene (LOC110373801/ XP_063898125.1) was employed as a query sequence for a BlastP search against the NCBI non-redundant (nr) protein sequences database. The top 100 hits were retrieved, and the resulting protein sequences were curated to remove duplicates and to retain only a single representative per species in cases of multiple hits from different genome assemblies, resulting in the exclusion of 13 genes.

The selected protein sequences were aligned using the MUSCLE tool (version 3.8.425) online, employing default settings. This alignment was then used to construct a phylogenetic tree using the IQ-TREE2 web server (Minh et al., 2020; Trifinopoulos et al., 2016). The tree was built with 1000 bootstrap replications, and the best-fit model (WAG+R5) was selected based on the Bayesian Information Criterion. One sequence (XP_013177522.1/PREDICTED: uncharacterized protein LOC106124990 from *Papilio xuthus*) was excluded from the analysis due to its having over 50% gaps compared to other sequences. Ultimately, 86 of the initial 100 protein sequences were included in the final tree analysis.

Visualization of the phylogenetic tree was conducted using the ggtree package (Yu et al., 2017), with bootstrap support values annotated on each node. The multiple sequence alignment file, used as input, is provided with amino acid residues coloured by type. Taxonomic classifications for each species were extracted from the NCBI Taxonomy database using a custom Python script.

### CRISPR-Cas9 knockout of the *HaVipR1* gene

The wild-type strain SCD that was originally collected from Côte D’Ivoire (Ivory Coast, Africa) in the 1970s was kindly provided by Bayer Crop Science in 2001 (Yang et al., 2009). This strain has been maintained in the laboratory without exposure to insecticides or Bt toxins for over 30 years, and it is susceptible to both Bt toxins and chemical insecticides (Yang et al., 2009). The *HaVipR1* gene (LOC110373801; HaOG214548) of the SCD strain was knocked out with the CRISPR/Cas9 genome editing tool (Wang et al., 2016) to produce a knockout strain homozygous for a 16-bp deletion in exon 3 of HaVip3R1.

### Toxicity of Vip3Aa to SCD and the knockout strain were determined with diet overlay bioassays

The Vip3Aa protoxin was provided by the Institute of Plant Protection, Chinese Academy of Agricultural Sciences (CAAS), Beijing, China. Protoxin in solution was prepared by diluting the stock suspensions with a 0.01 M, pH 7.4, phosphate buffer solution (PBS). Liquid artificial diet (1000 µl) was dispensed into each well (surface area = 2 cm^2^) of a 24-well plate. After the diet cooled and solidified, 100 µl of the Vip3Aa protein solution was applied evenly to the diet surface in each well. A single unfed neonate (24 h old) was put in each well after the Bt protein solution had dried at room temperature. Mortality was recorded after 7 days. Larvae were considered as dead if they died or weighed less than 5 mg at the end of bioassays.

The sgRNA target sequences (5’-GGCACCACGATAGGAAGATGTGG-3’, underlined is the PAM sequence) were selected at exon 3 of *HaVipR1* according to the principle of 5’-GGN18NGG-3’ (Fig 3). The template DNA was made with PCR-based fusion of two oligonucleotides: one is the specific oligonucleotide(CRISPR-F) encoding T7 polymerase-binding site and the sgRNA target sequence GGN18, and the other is the universal oligonucleotide(sgRNA-R) encoding the remaining sgRNA sequences. The fusion PCR reaction mixture (50 µl) consisted of 25 µl of PrimeSTAR® HS (Premix) (TaKaRa, Dalian, China), 2 µl of 10 µM CRISPR-F, 2 µl of 10 µM sgRNA-R, and 21 µl of ddH2O. PCR was performed at 98 ^°^C for 30 s, 35 cycles of (98 °C for 10 s, 60 °C for 20 s, 72 °C for 30 s), followed by 72 °C for 10 min, and held at 12 °C. The PCR products were then purified (using the QIAquick® PCR Purification Kit, (QIAGEN, Hilden, Germany). The sgRNA was synthesized by in vitro transcription utilizing the GeneArt™ Precision gRNA Synthesis Kit (Thermo Fisher Scientific, Shanghai, China) according to the manufacturer’s instruction. The Cas9 protein (GeneArt™ Platinum™ Cas9 Nuclease) were purchased from Thermo Fisher Scientific (Shanghai, China).

The collection and preparation of eggs were carried out as previously reported (Wang et al., 2016). Briefly, fresh eggs laid within 2 hours were washed down from the gauzes using a 1% sodium hypochlorite solution and rinsed with distilled water. After suction filtration, the eggs were lined up on a microscope slide fixed with double-sided adhesive tape. Approximately one nano-liter mix of sgRNA (300 ng/μl) and Cas9 protein (100 ng/μl) were injected into individual eggs using a FemtoJet and InjectMan NI 2 microinjection system (Eppendorf, Hamburg, Germany). The microinjection process was completed within 2 hours. Injected eggs were then placed at 26 ± 1 °C, 60 ± 10% RH for hatching.

To identify the indel mutations at exon 3 of *HaVipR1*, a pair of specific primers (forward: 5’-AACGTGAGGTTACTCATGCCTT-3’, reverse: 5’-GCCAAGCTTATTTTGACCAGCC-3’) was designed to amplify a ∼500-bp fragment flanking exon 3. The PCR fragments were amplified from genomic DNA samples of individual insects, and then sent to Life Technology (Shanghai, China) for direct sequencing, using the forward primer used as the sequencing primer. To determine the association between the Vip3Aa resistance phenotype and the 16-bp deletion mutation of *HaVipR1*, we performed a chi-square test of independence. Observed genotype frequencies for both treated (survivors) and untreated groups (control) were compared to expected frequencies under the assumption of genotype and treatment independence. Expected frequencies were calculated based on the total counts of each genotype across both groups.

## Acknowledgments

We gratefully acknowledge the support received from various organizations and funding bodies. Ashley Tessnow was supported by a travel grant (CLW1602) from the Cotton Research & Development Corporation (CRDC) and (16-413) from Cotton Incorporated. Craig J. Anderson was funded by the Commonwealth Scientific and Industrial Research Organisation (CSIRO) for his Postdoctoral research. This work was supported by several projects from the CRDC, including CSE1201 (original), CSE112, CSE104C, CSE0002, and CSE1103. We appreciate the contributions of these organizations to our research.

## References

Bel, Y., Jakubowska, A. K., Costa, J., Herrero, S., & Escriche, B. (2013). Comprehensive Analysis of Gene Expression Profiles of the Beet Armyworm Spodoptera exigua Larvae Challenged with Bacillus thuringiensis Vip3Aa Toxin. PLoS ONE, 8(12), e81927. 10.1371/journal.pone.0081927

Bergamasco, V. B., Mendes, D. R., Fernandes, O. A., Desidério, J. A., & Lemos, M. V. (2013). Bacillus thuringiensis Cry1Ia10 and Vip3Aa protein interactions and their toxicity in Spodoptera spp. (Lepidoptera). J Invertebr Pathol, 112(2), 152–158. 10.1016/j.jip.2012.11.011

Bernardi, O., Bernardi, D., Amado, D., Sousa, R. S., Fatoretto, J., Medeiros, F. C. L., Conville, J., Burd, T., & Omoto, C. (2015). Resistance Risk Assessment of Spodoptera frugiperda (Lepidoptera: Noctuidae) and Diatraea saccharalis (Lepidoptera: Crambidae) to Vip3Aa20 Insecticidal Protein Expressed in Corn. Journal of Economic Entomology, 108(6), 2711–2719. 10.1093/jee/tov219

Bernardi, O., Bernardi, D., Horikoshi, R. J., Okuma, D. M., Miraldo, L. L., Fatoretto, J., Medeiros, F. C., Burd, T., & Omoto, C. (2016). Selection and characterization of resistance to the Vip3Aa20 protein from Bacillus thuringiensis in Spodoptera frugiperda. Pest Manag Sci, 72(9), 1794–1802. 10.1002/ps.4223

Castagnola, A., & Jurat-Fuentes, J. L. (2016). Intestinal regeneration as an insect resistance mechanism to entomopathogenic bacteria. Current Opinion in Insect Science, 15, 104–110. 10.1016/j.cois.2016.04.008

Castro-Gomes, T., Corrotte, M., Tam, C., & Andrews, N. W. (2016). Plasma Membrane Repair Is Regulated Extracellularly by Proteases Released from Lysosomes. PLoS ONE, 11(3), e0152583. 10.1371/journal.pone.0152583

Catchen, J., Hohenlohe, P. A., Bassham, S., Amores, A., & Cresko, W. A. (2013). Stacks: an analysis tool set for population genomics. Mol Ecol, 22(11), 3124–3140. 10.1111/mec.12354

Chakroun, M., Banyuls, N., Bel, Y., Escriche, B., & Ferré, J. (2016). Bacterial Vegetative Insecticidal Proteins (Vip) from Entomopathogenic Bacteria. Microbiol Mol Biol Rev, 80(2), 329–350. 10.1128/mmbr.00060-15

Chakroun, M., Banyuls, N., Walsh, T., Downes, S., James, B., & Ferré, J. (2016). Characterization of the resistance to Vip3Aa in Helicoverpa armigera from Australia and the role of midgut processing and receptor binding. Sci Rep, 6, 24311. 10.1038/srep24311

Gomis-Cebolla, J., Wang, Y., Quan, Y., He, K., Walsh, T., James, B., Downes, S., Kain, W., Wang, P., Leonard, K., Morgan, T., Oppert, B., & Ferre, J. (2018). Analysis of cross-resistance to Vip3 proteins in eight insect colonies, from four insect species, selected for resistance to Bacillus thuringiensis insecticidal proteins. J Invertebr Pathol, 155, 64–70. 10.1016/j.jip.2018.05.004

Gomis-Cebolla, J., Wang, Y., Quan, Y., He, K., Walsh, T., James, B., Downes, S., Kain, W., Wang, P., Leonard, K., Morgan, T., Oppert, B., & Ferré, J. (2018). Analysis of cross-resistance to Vip3 proteins in eight insect colonies, from four insect species, selected for resistance to Bacillus thuringiensis insecticidal proteins. Journal of Invertebrate Pathology, 155, 64–70. 10.1016/j.jip.2018.05.004

Gupta, M., Kumar, H., & Kaur, S. (2021). Vegetative Insecticidal Protein (Vip): A Potential Contender From Bacillus thuringiensis for Efficient Management of Various Detrimental Agricultural Pests [Review]. Frontiers in Microbiology, 12. 10.3389/fmicb.2021.659736

Hernández-Martínez, P., Gomis-Cebolla, J., Ferré, J., & Escriche, B. (2017). Changes in gene expression and apoptotic response in Spodoptera exigua larvae exposed to sublethal concentrations of Vip3 insecticidal proteins. Sci Rep, 7(1), 16245. 10.1038/s41598-017-16406-1

Hou, X., Han, L., An, B., & Cai, J. (2021). Autophagy induced by Vip3Aa has a pro-survival role in Spodoptera frugiperda Sf9 cells. Virulence, 12(1), 509–519. 10.1080/21505594.2021.1878747

International Service for the Acquisition of Agri-biotech, A. (2024). GM Approval Database - Events List [Accessed: 2024-06-04]. https://www.isaaa.org/gmapprovaldatabase/eventslist/default.asp

Jakubowska, A. K., Caccia, S., Gordon, K. H., Ferré, J., & Herrero, S. (2010). Downregulation of a chitin deacetylase-like protein in response to baculovirus infection and its application for improving baculovirus infectivity. J Virol, 84(5), 2547–2555. 10.1128/jvi.01860-09

Jiang, K., Mei, S. Q., Wang, T. T., Pan, J. H., Chen, Y. H., & Cai, J. (2016). Vip3Aa induces apoptosis in cultured Spodoptera frugiperda (Sf9) cells. Toxicon, 120, 49–56. 10.1016/j.toxicon.2016.07.019

Jin, M., Shan, Y., Peng, Y., Wang, W., Zhang, H., Liu, K., Heckel, D. G., Wu, K., Tabashnik, B. E., & Xiao, Y. (2023). Downregulation of a transcription factor associated with resistance to Bt toxin Vip3Aa in the invasive fall armyworm. Proceedings of the National Academy of Sciences, 120(44), e2306932120. doi:10.1073/pnas.2306932120

Jurat-Fuentes, J. L., Heckel, D. G., & Ferré, J. (2021). Mechanisms of Resistance to Insecticidal Proteins from Bacillus thuringiensis. Annual Review of Entomology, 66(1), 121–140. 10.1146/annurev-ento-052620-073348

Kahn, T. W., Chakroun, M., Williams, J., Walsh, T., James, B., Monserrate, J., & Ferré, J. (2018). Efficacy and Resistance Management Potential of a Modified Vip3C Protein for Control of Spodoptera frugiperda in Maize. Scientific Reports, 8(1), 16204. 10.1038/s41598-018-34214-z

Kerns, D. D., Yang, F., Kerns, D. L., Stewart, S. D., & Jurat-Fuentes, J. L. (2023). Reduced binding associated with resistance to Vip3Aa in the corn earworm, Helicoverpa zea. bioRxiv, 2023.2007.2007.548161. 10.1101/2023.07.07.548161

Knight, K. M., Head, G. P., & Rogers, D. J. (2021). Successful development and implementation of a practical proactive resistance management plan for Bt cotton in Australia. Pest Management Science, 77(10), 4262–4273. 10.1002/ps.6490

Kurtz, R. W., McCaffery, A., & O’Reilly, D. (2007). Insect resistance management for Syngenta’s VipCot transgenic cotton. J Invertebr Pathol, 95(3), 227–230. 10.1016/j.jip.2007.03.014

Langmead, B., & Salzberg, S. L. (2012). Fast gapped-read alignment with Bowtie 2. Nat Methods, 9(4), 357–359. 10.1038/nmeth.1923

Lee, M. K., Walters, F. S., Hart, H., Palekar, N., & Chen, J. S. (2003). The mode of action of the Bacillus thuringiensis vegetative insecticidal protein Vip3A differs from that of Cry1Ab delta-endotoxin. Appl Environ Microbiol, 69(8), 4648–4657. 10.1128/aem.69.8.4648-4657.2003

Li, H. (2018). Minimap2: pairwise alignment for nucleotide sequences. Bioinformatics, 34(18), 3094–3100. 10.1093/bioinformatics/bty191

Lin, S., Oyediran, I., Niu, Y., Brown, S., Cook, D., Ni, X., Zhang, Y., Reay-Jones, F. P. F., Chen, J. S., Wen, Z., Dimase, M., & Huang, F. (2022). Resistance Allele Frequency to Cry1Ab and Vip3Aa20 in Helicoverpa zea (Boddie) (Lepidoptera: Noctuidae) in Louisiana and Three Other Southeastern U.S. States. Toxins, 14(4), 270. https://www.mdpi.com/2072-6651/14/4/270

Liu, J. G., Yang, A. Z., Shen, X. H., Hua, B. G., & Shi, G. L. (2011). Specific binding of activated Vip3Aa10 to Helicoverpa armigera brush border membrane vesicles results in pore formation. J Invertebr Pathol, 108(2), 92–97. 10.1016/j.jip.2011.07.007

Liu, Z., Liao, C., Zou, L., Jin, M., Shan, Y., Quan, Y., Yao, H., Zhang, L., Wang, P., Liu, Z., Wang, N., Li, A., Liu, K., Heckel, D. G., Wu, K., & Xiao, Y. (2024). Retrotransposon-mediated variation of a chitin synthase gene confers insect resistance to *Bacillus thuringiensis* Vip3Aa toxin. bioRxiv, 2024.2002.2018.580648. 10.1101/2024.02.18.580648

Love, M. I., Huber, W., & Anders, S. (2014). Moderated estimation of fold change and dispersion for RNA-seq data with DESeq2. Genome Biology, 15(12), 550. 10.1186/s13059-014-0550-8

Mahon, R. J., Downes, S. J., & James, B. (2012). Vip3A Resistance Alleles Exist at High Levels in Australian Targets before Release of Cotton Expressing This Toxin. PLoS ONE, 7(6), e39192. 10.1371/journal.pone.0039192

Mahon, R. J., & Young, S. (2010). Selection Experiments to Assess Fitness Costs Associated With Cry2Ab Resistance in Helicoverpa armigera (Lepidoptera: Noctuidae). Journal of Economic Entomology, 103(3),835–842. 10.1603/ec09330

McNeil, P. L., & Kirchhausen, T. (2005). An emergency response team for membrane repair. Nature Reviews Molecular Cell Biology, 6(6), 499–505. 10.1038/nrm1665

Mihelič, M., & Turk, D. (2007). Two decades of thyroglobulin type-1 domain research. Biological Chemistry, 388(11), 1123–1130. doi:10.1515/BC.2007.155

Pearce, S. L., Clarke, D. F., East, P. D., Elfekih, S., Gordon, K. H. J., Jermiin, L. S., Mcgaughran, A., Oakeshott, J. G., Papanikolaou, A., Perera, O. P., Rane, R. V., Richards, S., Tay, W. T., Walsh, T. K., Anderson, A., Anderson, C. J., Asgari, S., Board, P. G., Bretschneider, A., … Wu, Y. D. (2017). Genomic innovations, transcriptional plasticity and gene loss underlying the evolution and divergence of two highly polyphagous and invasive Helicoverpa pest species. BMC Biology, 15(1). 10.1186/s12915-017-0402-6

Pezzini, D., Taylor, K. L., Reisig, D. D., & Fritz, M. L. (2024). Cross-pollination in seed-blended refuge and selection for Vip3A resistance in a lepidopteran pest as detected by genomic monitoring. Proceedings of the National Academy of Sciences, 121(13), e2319838121. doi:10.1073/pnas.2319838121

Pickett, B. R., Gulzar, A., Ferre, J., & Wright, D. J. (2017). Bacillus thuringiensis Vip3Aa Toxin Resistance in Heliothis virescens (Lepidoptera: Noctuidae). Appl Environ Microbiol, 83(9). 10.1128/AEM.03506-16

Pigott, C. R., & Ellar, D. J. (2007). Role of receptors in Bacillus thuringiensis crystal toxin activity. Microbiol Mol Biol Rev, 71(2), 255–281. 10.1128/mmbr.00034-06

Ruiz de Escudero, I., Banyuls, N., Bel, Y., Maeztu, M., Escriche, B., Muñoz, D., Caballero, P., & Ferré, J. (2014). A screening of five Bacillus thuringiensis Vip3A proteins for their activity against lepidopteran pests. J Invertebr Pathol, 117, 51–55. 10.1016/j.jip.2014.01.006

Saikhedkar, N., Summanwar, A., Joshi, R., & Giri, A. (2015). Cathepsins of lepidopteran insects: Aspects and prospects. Insect Biochemistry and Molecular Biology, 64, 51–59. 10.1016/j.ibmb.2015.07.005

Tabashnik, B. E., & Carrière, Y. (2020). Evaluating Cross-resistance Between Vip and Cry Toxins of Bacillus thuringiensis. J Econ Entomol, 113(2), 553–561. 10.1093/jee/toz308

Tabashnik, B. E., Fabrick, J. A., & Carrière, Y. (2023). Global Patterns of Insect Resistance to Transgenic Bt Crops: The First 25 Years. Journal of Economic Entomology, 116(2), 297–309. 10.1093/jee/toac183

Tay, W. T., Mahon, R. J., Heckel, D. G., Walsh, T. K., Downes, S., James, W. J., Lee, S.-F., Reineke, A., Williams, A. K., & Gordon, K. H. J. (2015). Insect Resistance to Bacillus thuringiensis Toxin Cry2Ab Is Conferred by Mutations in an ABC Transporter Subfamily A Protein. PLOS Genetics, 11(11), e1005534. 10.1371/journal.pgen.1005534

Turk, V., Stoka, V., Vasiljeva, O., Renko, M., Sun, T., Turk, B., & Turk, D. (2012). Cysteine cathepsins: from structure, function and regulation to new frontiers. Biochim Biophys Acta, 1824(1), 68–88. 10.1016/j.bbapap.2011.10.002

United States Environmental Protection, A. (2024). Current and Previously Registered Section 3 Plant-Incorporated Protectants [Accessed: 2024-06-04]. https://www.epa.gov/ingredients-used-pesticide-products/current-and-previously-registered-section-3-plant-incorporated

Vidak, E., Javoršek, U., Vizovišek, M., & Turk, B. (2019). Cysteine Cathepsins and their Extracellular Roles: Shaping the Microenvironment. Cells, 8(3). 10.3390/cells8030264

Walsh, T. K., Downes, S. J., Gascoyne, J., James, W., Parker, T., Armstrong, J., & Mahon, R. J. (2014). Dual Cry2Ab and Vip3A resistant strains of Helicoverpa armigera and Helicoverpa punctigera (Lepidoptera: Noctuidae); testing linkage between loci and monitoring of allele frequencies. J Econ Entomol, 107(4), 1610–1617. 10.1603/ec13558

Wells, J. N., & Feschotte, C. (2020). A Field Guide to Eukaryotic Transposable Elements. Annu Rev Genet, 54, 539–561. 10.1146/annurev-genet-040620-022145

Wen, Z., Conville, J., Matthews, P., Hootman, T., Himes, J., Wong, S., Huang, F., Ni, X., Chen, J. S., & Bramlett, M. (2023). More than 10 years after commercialization, Vip3A-expressing MIR162 remains highly efficacious in controlling major Lepidopteran maize pests: laboratory resistance selection versus field reality. Pesticide Biochemistry and Physiology, 192, 105385. 10.1016/j.pestbp.2023.105385

Yang, F., Kerns, D. L., Little, N. S., Santiago González, J. C., & Tabashnik, B. E. (2021). Early Warning of Resistance to Bt Toxin Vip3Aa in Helicoverpa zea. Toxins (Basel), 13(9). 10.3390/toxins13090618

Yang, F., Santiago González, J. C., Sword, G. A., & Kerns, D. L. (2021). Genetic basis of resistance to the Vip3Aa Bt protein in Helicoverpa zea. Pest Manag Sci, 77(3), 1530–1535. 10.1002/ps.6176

Yu, G., Smith, D. K., Zhu, H., Guan, Y., & Lam, T. T.-Y. (2017). ggtree: an r package for visualization and annotation of phylogenetic trees with their covariates and other associated data. Methods in Ecology and Evolution, 8(1), 28–36. 10.1111/2041-210X.12628

Zhang, J., Zhang, F., Tay, W. T., Robin, C., Shi, Y., Guan, F., Yang, Y., & Wu, Y. (2022). Population genomics provides insights into lineage divergence and local adaptation within the cotton bollworm. Mol Ecol Resour, 22(5), 1875–1891. 10.1111/1755-0998.13581

