## Supplementary Material and Information for "Identification of a novel resistance gene which provides insight into Vip3Aa mode of action in *Helicoverpa armigera*"

**This PDF file includes:**

Supporting text
Figures S1 to S7
Tables S1 to S4
SI References

**Supporting Information Text**

**Methods**

**Analysis of expression of HaVipR1 homologue in *Spodoptera frugiperda* mid-gut and Sf9 cell lines.**
The Short-Read Archive (SRA) browser was used to find RNA sequence data from either Sf9 cell lines or *S. frugiperda* mid-gut derived tissues. Seven mid-gut samples and 13 Sf9 samples (Table S4) were downloaded and aligned to the *S. frugiperda* reference using Hisat2 with default parameters (GCF_023101765.2, "AGI-APGP_CSIRO_Sfru_2.0"). The homologue for HaVipR1 in *S. frugiperda* was found using mmseqs easy-rbh and was identified as LOC118272819 "thyroglobulin" gene. Overall gene expression was analysed across samples using DeSeq2 and the normalised read counts for the Sf *HaVipR1* homologue was derived using plotCounts.

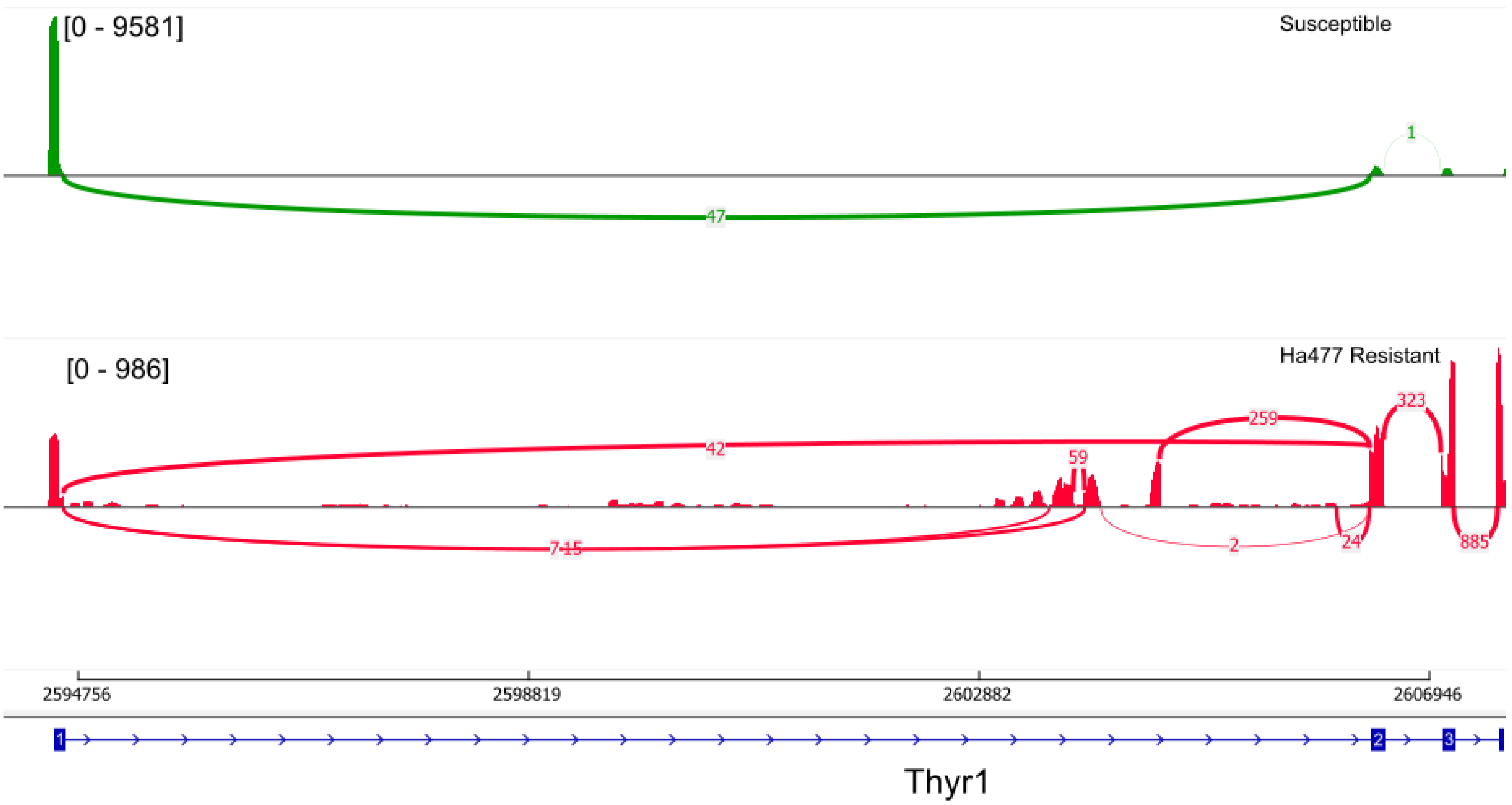


Fig. S1. **Sashimi plot demonstrating differential splicing of the *HaVipR1* transcript in the Ha477 resistant line**. Three samples from the susceptible and resistant (Ha477) H. armigera lines assessed were mapped to the de novo polished assembly from an individual from the Ha477 resistant line. The *HaVipR1* gene structure was identified using miniport and is annotated to show the first three coding exons. The resistant line displays splicing present near the 3' end of the repeat rich region with direct links to the second exon. No aberrant splicing is identified in the susceptible line.

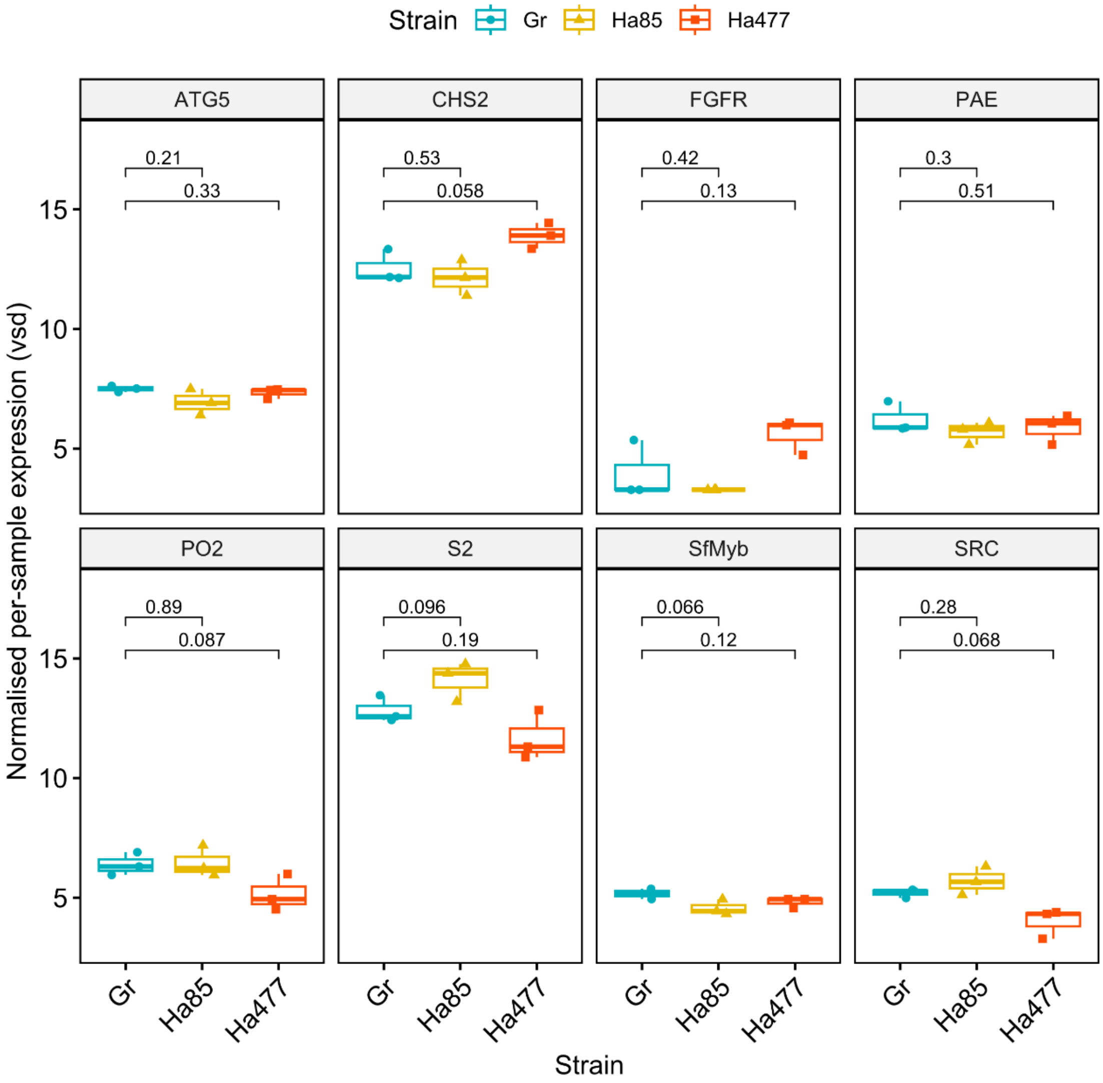


**Fig. S2. Analysis of expression of 8 other Vip3A related genes from *S. frugiperda* in pooled mid-gut transcriptome data from the susceptible and two resistant allelic (Ha85 and Ha477) lines of *H. armigera*.** Variance stabilised expression values are shown for each strain and statistical significance calculated using Students t-test for each of the resistant lines in comparison to the susceptible line. p-values are provided for each comparison.

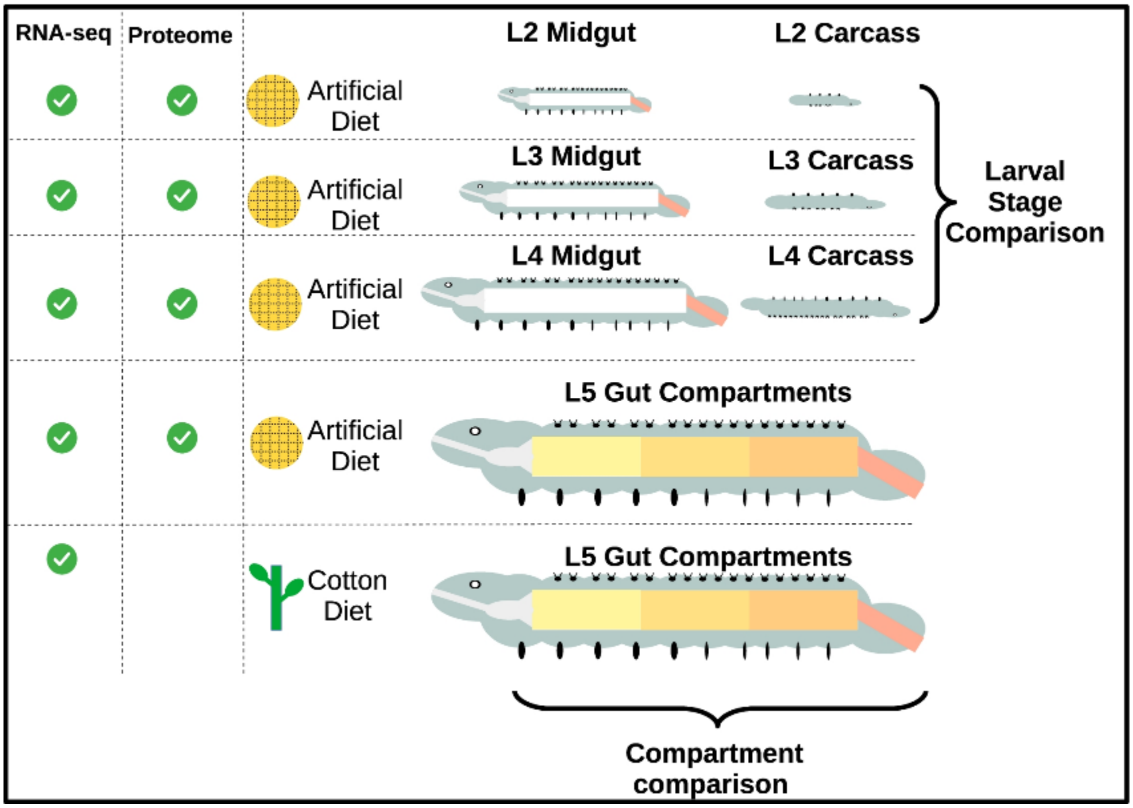


**Fig. S3. Descriptions of sample tissue for transcriptomic profiling of H. armigera, reproduced from (Ioannidis et al., 2022)**. Whole insects were used for stages L2, L3 and L4 and either the midgut (MG) or carcass (C) analysed. For the L5 larvae, guts were dissected out and separated into 5 compartments corresponding to the foregut (FG), anterior midgut (AMG), middle midgut (MMG), posterior midgut (PMG), and hindgut (HG). Four biological replicates from each gut condition were included in the analysis and each biological replicated consisted of at least five tissues. Type or paste legend here. Paste figure above the legend.

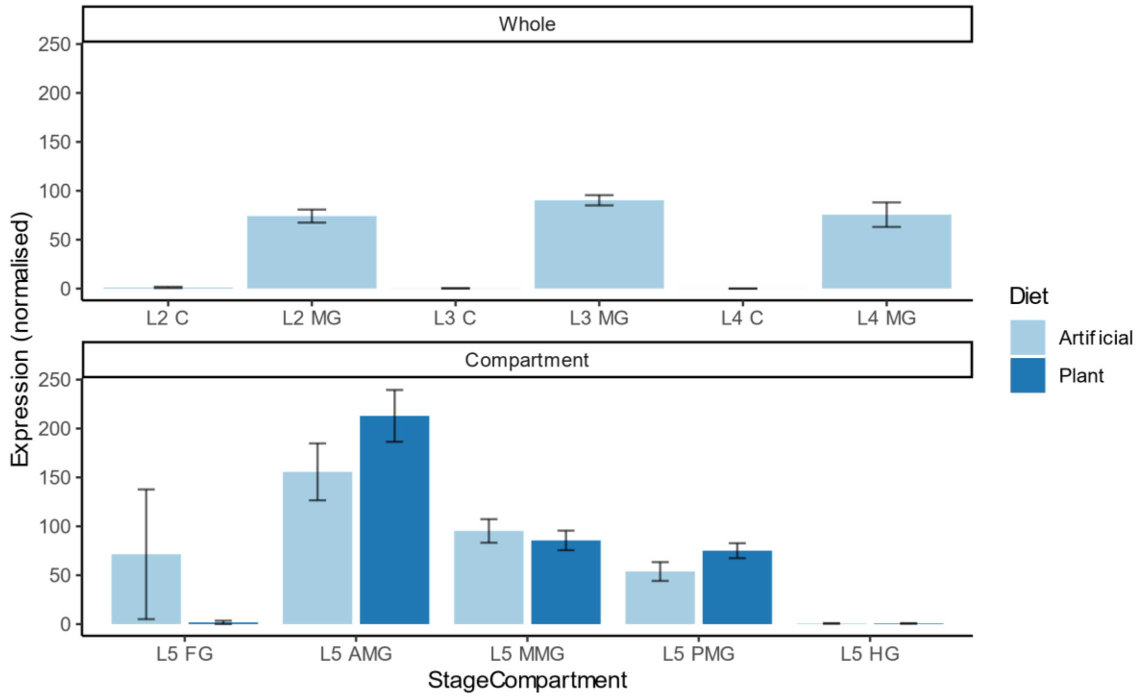


**Fig. S4. Expression values for HaVipR1 in the spatio-transcriptome dataset generated from (Ioannidis et al., 2022).** Results for the whole body stages (L2, L3 and L4) and the compartment analysis (L5) have been shown along with the reported standard error.

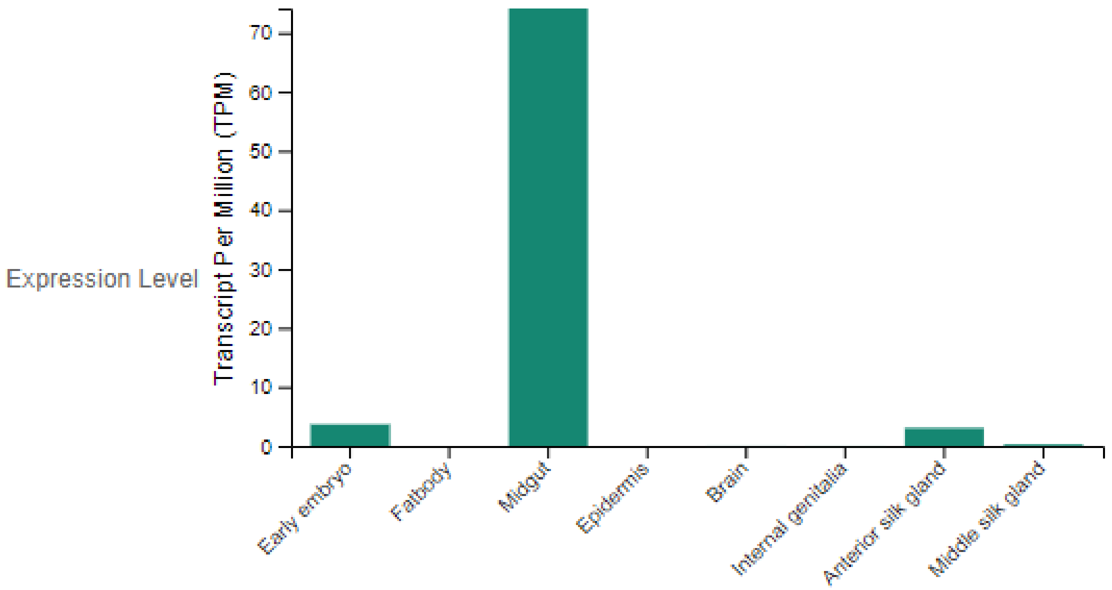


**Fig. S5. Expression of the homologue of HaVipR1 in *Bombyx mori* indicates peak of expression of the gene in the midgut.** The homologous gene for Bombyx mori was found to be LOC101735440 according to the phylogeny and this was used to search the SilkWorm database (Kawamoto & Katsuma, 2022). The matching record in the database is accession KWMTBOMO00841. The record can be accessed at this address https://silkbase.ab.a.u-tokyo.ac.jp/cgi-bin/entryview_gm2.cgi?clone_name=%20KWMTBOMO00841 .

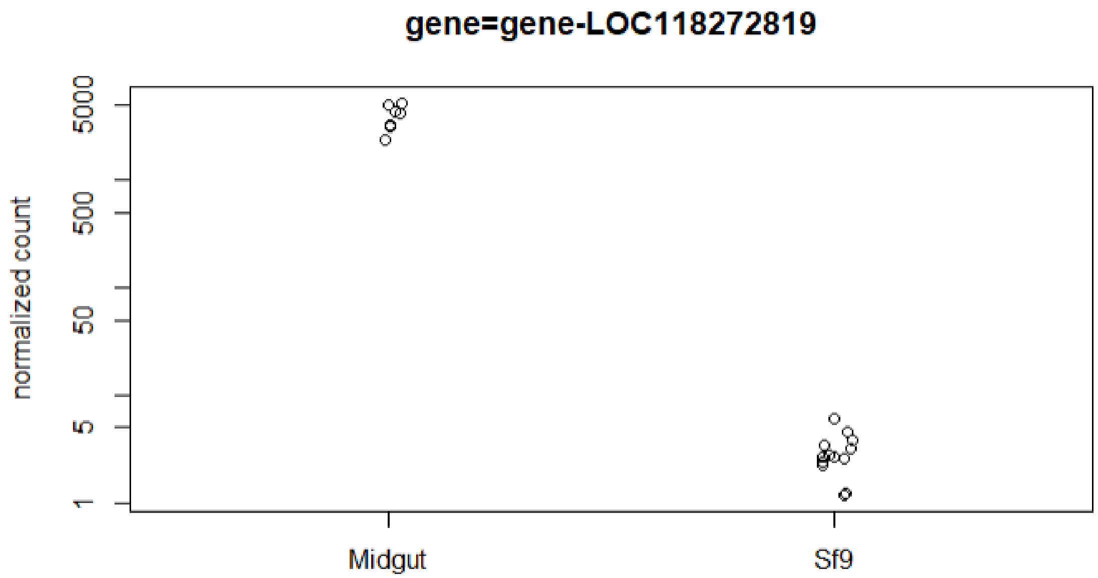


**Fig. S6. Expression of HaVipR1 homolog in Spodoptera frugiperda mid-gut tissues and Sf9 cell lines.** Samples analysed are described in Table S4. Individual samples were aligned to the S. frugiperda reference and the normalised read counts for the homolog to HaVipR1 was extracted from DeSeq2.

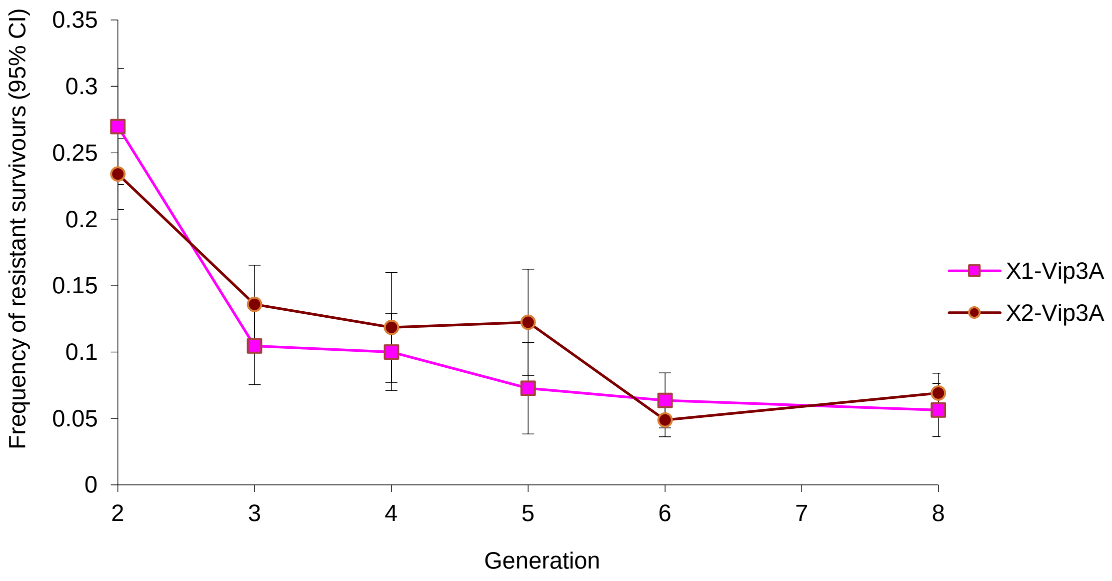


**Fig. S7. In the absence of selection the HaVipR1 phenotype decreases over time.** The Vip3A resistant line Ha85 was maintained in the laboratory in the absence of selection for 8 generations and for both replicates (X1 and X2) the frequency of the Vip3A resistance phenotype declined over time.

**Table S1. Number of RadTag loci per chromosome in female informative cross.**

| Chromosome | Chromosome identifier | Number RadTag loci |
|---|---|---|
| *Z* | NC_064776.1 | 2,117 |
| *1* | NC_064777.1 | 423 |
| *2* | NC_064778.1 | 1,150 |
| *3* | NC_064779.1 | 1,094 |
| *4* | NC_064780.1 | 1,427 |
| *5* | NC_064781.1 | 1,250 |
| *6* | NC_064782.1 | 1,006 |
| *7* | NC_064783.1 | 1,237 |
| *8* | NC_064784.1 | 1,195 |
| *9* | NC_064785.1 | 1,451 |
| *10* | NC_064786.1 | 496 |
| *11* | NC_064787.1 | 1,289 |
| *12* | NC_064788.1 | 1,181 |
| *13* | NC_064789.1 | 1,021 |
| *14* | NC_064790.1 | 1,756 |
| *15* | NC_064791.1 | 1,072 |
| *16* | NC_064792.1 | 1,176 |
| *17* | NC_064793.1 | 1,188 |
| *18* | NC_064794.1 | 1,046 |
| *19* | NC_064795.1 | 866 |
| *20* | NC_064796.1 | 1,178 |
| *21* | NC_064797.1 | 1,005 |
| *22* | NC_064798.1 | 1,159 |
| *23* | NC_064799.1 | 530 |
| *24* | NC_064800.1 | 1,203 |
| *25* | NC_064801.1 | 713 |
| *26* | NC_064802.1 | 746 |
| *27* | NC_064803.1 | 921 |
| *28* | NC_064804.1 | 1,021 |
| *29* | NC_064805.1 | 438 |
| *30* | NC_064806.1 | 565 |

**Table S2.** Mapping statistics for transcriptome samples from the two resistant (Ha85 and Ha477) and susceptible (GR) H. armigera lines.

| Strain | Sample | Total Reads (counts) | Reads mapped (counts) | Reads mapped (%) |
|---|---|---|---|---|
| **Ha85** | S36 | 24,158,222 | 17,180,488 | 71.12% |
| **Ha85** | S39 | 24,129,320 | 17,913,549 | 74.24% |
| **Ha85** | S43 | 24,586,196 | 18,030,907 | 73.34% |
| **Ha477** | S15 | 26,006,796 | 18,047,080 | 69.39% |
| **Ha477** | S1 | 20,265,480 | 14,324,331 | 70.68% |
| **Ha477** | S28 | 29,702,064 | 20,037,062 | 67.46% |
| **GR** | S44 | 24,018,042 | 17,388,181 | 72.40% |
| **GR** | S45 | 24,405,054 | 17,526,844 | 71.82% |
| **GR** | S46 | 24,205,510 | 17,930,612 | 74.08% |

**Table S3. Results from reciprocal best hit analysis of *S. frugiperda* (SF) Vip3A related genes and *H. armigera* (HA) genes.**

| SfGene | SfProt | SfGeneName | HaProt | HaGene | HaGeneName | Short name |
|---|---|---|---|---|---|---|
| LOC118275234 | XP_035449032.1 | Phenoloxidase subunit 2-like | XP_021182404.1 | LOC110370773 | phenoloxidase 1 | PO2 |
| LOC118279360 | XP_035454931.2 | Phenoloxidase-activating enzyme-like | XP_049706366.1 | LOC110371189 | phenoloxidase-activating enzyme | PAE |
| LOC118273105 | XP_050552797.1 | chitin synthase chs-2-like | XP_021184028.2 | LOC110371908 | chitin synthase chs-2-like | CHS2 |
| LOC118280749 | XP_035457001.2 | Scavenger receptor-C | XP_021184773.2 | LOC110372413 | MAM and LDL-receptor class A domain-containing protein 1 | SRC |
| LOC118277870 | XP_035452760.1 | Fibroblast growth factor receptor | XP_049693499.1 | LOC110373728 | fibroblast growth factor receptor homolog 1 | FGFR |
| LOC118262572 | XP_035429947.1 | Autophagy related gene 5 | XP_021187832.1 | LOC110374447 | autophagy protein 5 | ATG5 |
| LOC118263612 | XP_035431600.1 | Ribosomal S2 | XP_021191964.1 | LOC110377400 | 40S ribosomal protein S2 | S2 |
| LOC118279242 | XP_035454758.2 | myb protein | XP_049705840.1 | LOC126053489 | myb protein-like | SfMyb |

**Table S4. SRA identifiers from either Sf9 or *S. frugiperda* mid-gut samples analysed for HaVipR1 expression.**

| Sf9 | Midgut |
|---|---|
| SRR21707399 | SRR17042499 |
| SRR21707400 | SRR17042500 |
| SRR21707407 | SRR17042505 |
| SRR21707408 | SRR22019602 |
| SRR21707409 | SRR22019603 |
| SRR21707410 | SRR22019604 |
| SRR21707411 | SRR22019608 |
| SRR21707412 | |
| SRR21707413 | |
| SRR21707414 | |
| SRR21707415 | |
| SRR21707416 | |
| SRR21998727 | |

**SI References**

Ioannidis, P., Buer, B., Ilias, A., Kaforou, S., Aivaliotis, M., Orfanoudaki, G., Douris, V., Geibel, S., Vontas, J., & Denecke, S. (2022). A spatiotemporal atlas of the lepidopteran pest Helicoverpa armigera midgut provides insights into nutrient processing and pH regulation. *BMC Genomics*, *23*(1), 75. https://doi.org/10.1186/s12864-021-08274-x

Kawamoto, M., & Katsuma, S. (2022). SilkBase Update: Addition of Information on Predicted Protein 3D Structures. *Sanshi-Konchu Biotec*, *91*(3), 3_217-213_220. https://doi.org/10.11416/konchubiotec.91.3_217
